# Monkeypox virus spreads from cell-to-cell and leads to neuronal death in human neural organoids

**DOI:** 10.1101/2023.09.19.558432

**Authors:** Isabel Schultz-Pernice, Amal Fahmi, Francisco Brito, Matthias Liniger, Yen-Chi Chiu, Teodora David, Blandina I. Oliveira Esteves, Antoinette Golomingi, Beatrice Zumkehr, Markus Gerber, Damian Jandrasits, Roland Züst, Selina Steiner, Carlos Wotzkow, Fabian Blank, Olivier B. Engler, Artur Summerfield, Nicolas Ruggli, David Baud, Marco P. Alves

**Author notes:** Correspondence (M.P.A.).

## Abstract

In 2022-23, the world experienced the largest recorded monkeypox virus (MPXV) outbreak outside of endemic regions. Remarkably, cases of neurological manifestations were reported, some of which fatal. MPXV DNA and MPXV-specific antibodies were detected in the cerebrospinal fluid of encephalitis-affected patients, suggesting neuroinvasive potential of MPXV. We explored the susceptibility of neural tissue to MPXV infection using human neural organoids (hNOs) exposed to a primary isolate belonging to clade IIb lineage. The virus efficiently replicates in hNOs as indicated by the exponential increase of infectious viral loads and the elevated frequency of MPXV-positive cells over time. Electron microscopy imaging revealed the presence of viral particles as well as perinuclear viral factories. We observed susceptibility of several cell types to the virus, including neural progenitor cells and neurons. Furthermore, we detected the presence of viral antigens in neurites and in foci of grouped cells distributed throughout the tissue. In line with this, we documented significantly more cell-associated than released infectious virus, suggesting viral spread by cell-to-cell contact. Using an mNeonGreen-expressing recombinant MPXV, we confirmed cell-associated virus transmission through live-cell imaging. While hNOs displayed no evident outer morphological changes upon infection, we detected the formation of beads in neurites, a phenomenon commonly associated with neurodegenerative disorders. Live-cell imaging further confirmed the recurrent formation of neuritic beads in neurons in the days following MPXV infection, with bead formation preceding neurite-initiated cell death. Notably, treatment of MPXV infected hNOs with the antiviral drug tecovirimat resulted in a significant reduction of infectious viral loads by several orders of magnitude. Taken together, our findings suggest viral manipulation of axonal transport driving neuronal degeneration and identify a mechanism potentially contributing to MPXV-mediated neuropathology that may have therapeutic implications.

## INTRODUCTION

The first case of monkeypox virus (MPXV) infection without traceable contact to African population or fauna was registered in UK in early May 2022^1,2^. Since then, more than 100 countries reported cases of infection^1^, marking the largest recorded MPXV epidemic outside of African countries. During the 2022 outbreak, initial manifestations of mpox, the illness caused by MPXV, included fever, lethargy, myalgia, lymphadenopathy and headache, followed by skin lesion development^3–5^. While most clinical cases recorded during the epidemic were observed to be mild^3,5^, severe complications may develop. From 1985 to 2021, serious neurological manifestations, including confusion, seizures and encephalitis, have been recorded in around 3% of MPXV infected patients^6^, including three fatal cases^7,8^. Since May 2022, neurological manifestations have been reported in several patients affected by MPXV infection^9–20^, including two young, healthy men that died following the complications^9^. Animal studies have suggested neuroinvasive potential of MPXV, detecting viral DNA and/or infectious virus in the brain of several rodent species^21–33^ as well as in the central nervous system (CNS) of pregnant rhesus macaques and their fetuses^34^. This hypothesis is further supported by the detection of MPXV DNA in the cerebrospinal fluid (CSF) of one affected woman during the 2022 outbreak^11^ as well as the detection of intrathecally produced MPXV-specific antibodies in two cases^19,35^.

Despite first reports of deadly encephalitis cases dating back to 1987^7^, mechanisms driving acute neurological manifestations during MPXV infections have been poorly investigated in the human host. Possible causes include the previously restricted geographical distribution of the virus, as well as limited availability of material and accurate models to analyze the neuropathology in the human brain. In recent years, three-dimensional (3D) cell culture systems have emerged as a powerful tool to recapitulate processes shaping human organs in health and disease^36–39^. These *in vitro* systems have provided unprecedented opportunities to study host-pathogen interactions in the human host exploiting a complex system accessible to experimental manipulation^40–42^. Notably, human neural organoids (hNOs), also known as brain or cerebral organoids, represent a versatile *in vitro* system that recapitulates human brain development and provides an advanced model to study neurological disorders^43,44^. Motivated by the need to uncover the mechanisms underlying mpox-associated encephalitis, in this study, we investigated the consequences of MPXV neuroinvasion by assessing MPXV tropism and neural cell susceptibility to MPXV infection using hNOs. Our results indicate that MPXV leverages axonal transport for dissemination, leading to neuronal degeneration, and uncover a potential mechanism contributing to MPXV-induced neuropathology.

## RESULTS

### Human neural cells are susceptible to monkeypox virus infection

The generation of hNOs from human pluripotent stem cells, encompassing embryonic stem cells (ESCs, cell line H1) and induced pluripotent stem cells (iPSCs, cell line IMR90), was performed according to established methods (Supplementary Fig. 1a)^45,46^. To phenotype our cultures, hNO tissue of different developmental stages was sampled over a period of 112 days and processed for immunofluorescence characterization using well-established cell type markers to discriminate between class III β-tubulin (TUJ1)-expressing neurons^47–50^ and sex-determining region Y-box 2 (SOX2) positive neural progenitor cells (NPCs)^51,52^. During the first days of development, hNOs were observed encompassing high numbers of tightly clustered, small ventricular units, previously described as fluid-filled cavities^45^ surrounded by a compact but relatively thin sheath of SOX2-expressing NPCs (Fig. 1a, top panels). Isolated TUJ1-positive neurons were occasionally observed scattered around the ventricles (Fig. 1a, top panels). Throughout the first and second month of development, hNOs displayed increasing degrees of complexity, acquiring a typical layered organization^45^. By day 60 of tissue expansion, enlarged ventricles surrounded by several layers of tightly gathered NPCs became evident (Fig. 1a, central panels). Densely packed TUJ1-expressing neurons surrounded the ventricular units, forming a thick meshwork of somata and neurites (Fig. 1a, central panels). Finally, a nuclei-poor outer layer was observed on the organoids’ basal surface (Fig. 1a, central panels). As development progressed further, organoids were observed losing ventricular organization, displaying a scattered population of SOX2-positive cells nested within a neuron-dominated environment (Fig. 1a, bottom panels). Furthermore, astrocytes, characterized by glial fibrillary acidic protein (GFAP) expression^53^ in absence of SOX2 signal, were confirmed appearing in hNOs by the third month of maturation, in line with previous findings reporting the presence of astrocytes at low frequencies by day 45 of organoid development^54^.

**Figure 1:**
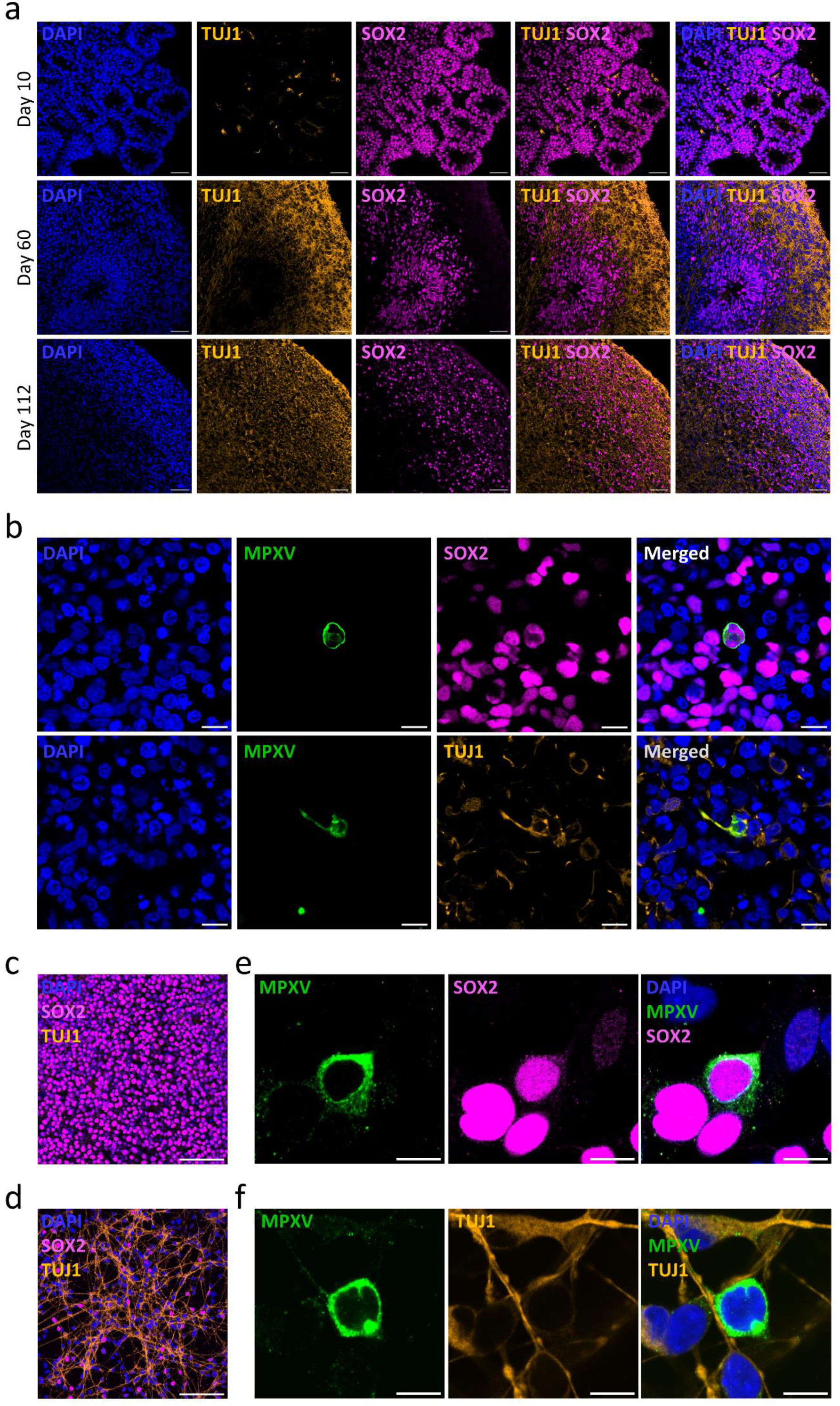
MPXV has a broad cellular tropism in hNOs and 2D neural cultures. (a) Representative micrographs showing the cell diversity and morphology of ESC-derived hNOs at selected developmental time points (10, 60, and 112 days). Selected markers of NPCs (SOX2, pink) and neurons (TUJ1, orange) are presented. DAPI, blue. Scale bar, 50 µm. (b) Representative micrographs illustrating the target cells of MPXV in 70-75 days old hNOs 10 days p.i. with a clade IIb lineage MPXV isolate at an MOI of 0.1 TCID_50_/cell. DAPI, blue; MPXV, green; SOX2, pink; TUJ1, orange. Scale bar, 10 µm. (c-d) Characterization of 2D NPC (c) and forebrain neuron (d) cultures. DAPI, blue; SOX2, pink; TUJ1, orange. Scale bar, 100 μm. (e-f) Exemplary image of 2D-cultured NPCs (e) and neurons (f) 8 hours p.i. with a clade IIb lineage MPXV isolate at an MOI of 0.01 TCID_50_/cell. DAPI, blue; MPXV, green; SOX2, pink; TUJ1, orange. Scale bar, 10 μm.

Aiming to shed light on the dynamics of MPXV replication within human neural tissue, we set out to analyze its tropism within our cultures. Eight independent batches of H1-derived hNOs and three distinct batches of IMR90-derived organoids of 70-110 days of culture were exposed to a 2022 primary isolate of MPXV belonging to clade IIb lineage. The infected hNOs were sampled at regular intervals and processed for immunofluorescence analysis (Supplementary Fig. 1b). In line with recent findings^55^, we observed viral antigen localizing to the cytoplasm of SOX2-expressing cells (Fig. 1b, top panels), indicating susceptibility of NPCs to MPXV infection. In contrast to previous reports^55^, hNO-enclosed neurons were observed to be susceptible to MPXV infection as well, as viral antigen was identified co-localizing with TUJ1 in both perikarya and neurites (Fig. 1b, bottom panels). Additionally, SOX2-negative but GFAP-expressing astrocytes were confirmed to display viral antigen signal within cell somata and extensions (Supplementary Fig. 2).

To further corroborate our findings in NPCs and neurons, the predominant cell types in hNOs, we established and subjected 2D NPC- and forebrain neuron-dominated cultures (henceforth addressed as “NPC cultures” and “forebrain neuron cultures”) to MPXV infection. High cell type purity of H1-derived NPC and forebrain neuron cultures was assessed through immunofluorescence staining with previously described markers (Fig. 1c and d). 2D NPC cultures showed characteristic teardrop cell morphology and rosette formation, along with a high density of SOX2-positive cells (Fig. 1c), confirming successful ESC transition to NPC state. Likewise, cells further differentiated towards a forebrain neuronal fate displayed robust neurite formation, encompassing dendrite and axon development, along with strong TUJ1 expression (Fig. 1d). In line with hNO-derived findings, immunofluorescence microscopy of NPC and forebrain neuron cultures confirmed presence of viral antigen in both cell types (Fig. 1e and f). As observed in hNO enclosed cells, MPXV signal was visualized within the cell body of both cell types, while fluorescent signal was additionally visible in cell extensions of 2D forebrain neurons (Fig. 1f).

### Human neural cells are permissive to monkeypox virus infection

We next set out to analyze the dynamics of MPXV replication by exposing hNOs to a low multiplicity of infection (MOI) of 0.1 TCID_50_/cell. Images, samples of tissue and supernatant were collected at regular intervals till 10 or 14 days post-infection (p.i.; Supplementary Fig. 1b). No evident alterations between mock-treated and MPXV-infected cultures were observed in outer morphology at any time point (Supplementary Fig. 3a and b). In addition, no significant or pronounced differences emerged through surface area analysis of H1-derived hNOs, with mock-treated organoids displaying an average surface area over all time points of 12.7 mm^2^, with an increase of 2.0 mm^2^ during the course of the whole experiment, compared to MPXV-challenged hNOs, that showed a mean size of 13.2 mm^2^ and 1.9 mm^2^ growth from day 0 to day 14 p.i. (mock vs. MPXV day 14 p.i., p = 0.8). Comparable, non-significant changes (mock vs. MPXV day 14 p.i., p = 0.2) in average surface area over all time points and surface area increase were determined for IMR90-derived organoids, despite hNOs displaying an overall smaller size compared to H1-derived cultures.

Viral replication and distribution in the tissue over time was investigated through immunofluorescence analysis. Viral particles were visualized using an antibody targeting the virus’s A27L protein, a conserved and multifunctional viral envelope protein known to mediate viral attachment and fusion to cell surfaces, as well as transport of the related vaccinia virus through microtubule interaction^56–59^. While no viral antigen was visible in mock-treated hNOs, small gatherings of MPXV-positive cells were detected in infected cultures as early as day 2 p.i. (Fig. 2a, Supplementary Fig. 3c and d). Viral antigen signal was subsequently observed steadily increasing over time, reaching a peak at day 10 p.i., at which numerous elliptical and elongated infected regions became evident around the organoids’ perimeter alongside scattered single cells reaching deep into the hNOs’ core (Fig. 2a, Supplementary Fig. 3c and d). Between day 10 and day 14 p.i., MPXV signal reached its maximum intensity, as no further increase could be detected (Fig. 2a, Supplementary Fig. 3c and d). Strikingly, MPXV antigen was observed consistently displaying a clustered distribution with positive cells arranged in delimited foci at the hNO’s surface layers (Fig. 2a and Supplementary Fig. 3c and d).

**Figure 2:**
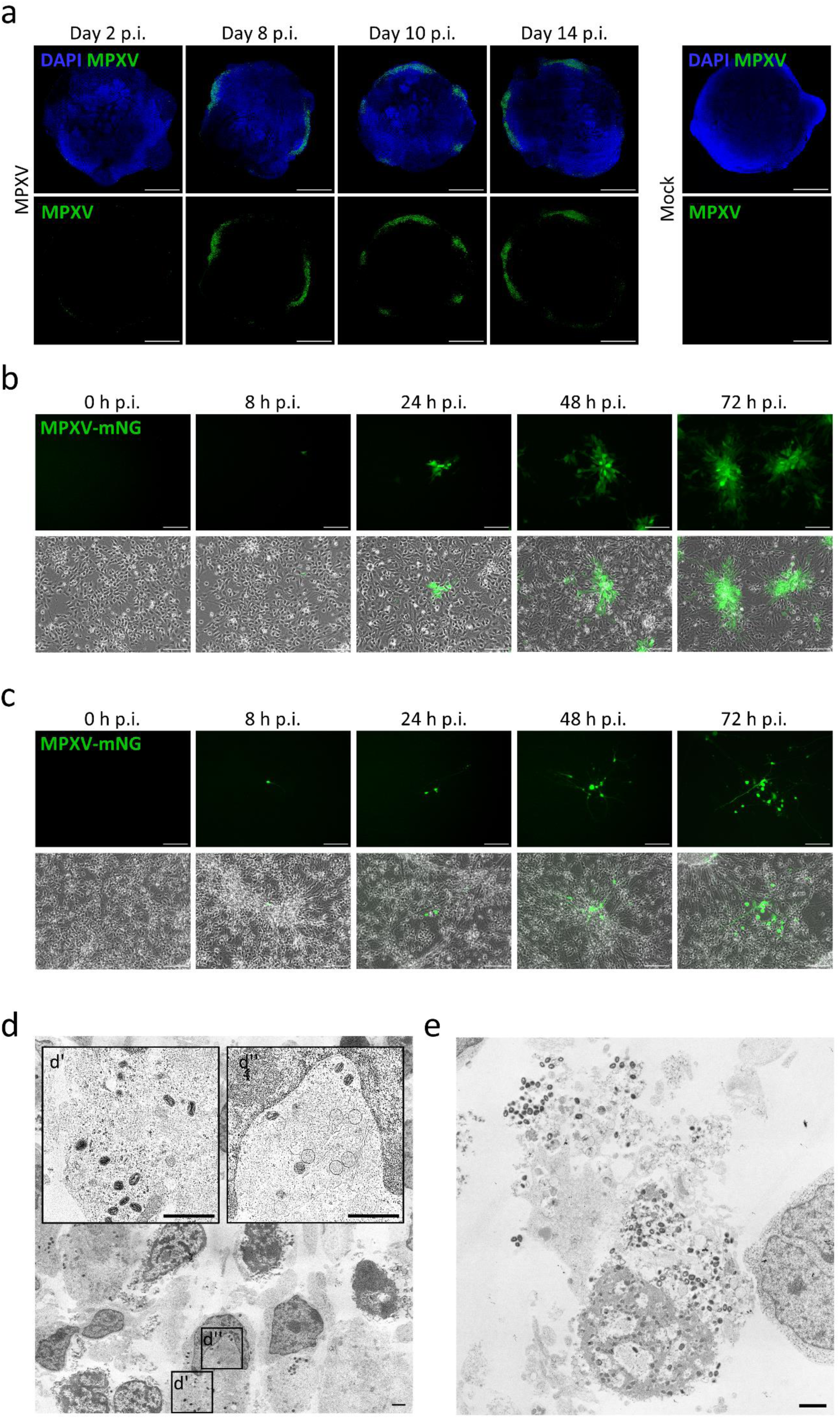
Productive MPXV infection in hNOs and 2D cultured neural cells. (a) Representative micrographs of hNOs infected with a clade IIb lineage MPXV isolate at a rate of 0.1 TCID_50_/cell at 83 days of development. Infection was followed by immunofluorescence from 2 to 14 days p.i. by whole-organoid imaging. DAPI, blue; MPXV, green. Scale bar, 1000 µm. (b) Representative micrographs depicting infection progression of 2D cultured NPCs with an mNG-expressing clade IIb lineage MPXV reporter strain at a rate of 0.01 TCID_50_/cell. Scale bar, 100 μm. (c) Representative images of 2D cultured forebrain neurons exposed with an mNG-expressing clade IIb lineage MPXV strain at a rate of 0.01 TCID_50_/cell. Scale bar, 100 μm. (d-e) Representative images acquired by transmission electron microscopy at 10 days p.i. of 95 days old hNOs infected with a clade IIb lineage MPXV isolate with an MOI of 0.1 TCID_50_/cell. (d’) Magnification demonstrating the presence of intracellular mature virions within cells. Scale bar, 1 µm. (d’’) Magnification showing the presence of viral particles at different stages of development alongside with perinuclear viral factories, displaying cytoskeletal condensations. Scale bar, 1 µm. (e) Illustration of degenerative signs, bursting and release of intracellular compartment containing high amounts of monkeypox virions. Scale bar, 1 µm.

Aiming to further analyze viral replication and impact on the predominant hNO cell populations, 2D NPC and forebrain neuron cultures were exposed to MPXV at an MOI of 0.01 TCID_50_/cell. As positive control, Vero E6 cells, highly susceptible to MPXV infection^60^, were challenged. Virus entry and replication, as well as cytopathic effect and distribution pattern within all cell types was investigated through microscopy analysis. Pronounced cell death was observed in Vero E6 cells as early as 48 hours p.i., with the appearance of multiple plaques that were documented to increase in size within the following two days (Supplementary Fig. 3e). In contrast, only a limited cytopathic effect of MPXV infection on NPCs and neurons was observed at 72-hour p.i. (Supplementary Fig. 3f and g). Seeking to further investigate virus spread in 2D neural cultures, we constructed an mNeonGreen (mNG)-expressing clade IIb lineage MPXV, derived from the previously applied viral isolate (Supplementary Fig. 3h). NPCs and forebrain neurons were subsequently exposed to the MPXV-mNG virus at an MOI of 0.01 TCID_50_/cell and imaged at defined time points. In both NPC and neuron cultures, scattered single cells were documented expressing mNG signal at 8 hours p.i. (Fig. 2b and c), a time point at which late stages of viral replication are expected to occur in susceptible cells and thereby indicative of successful virus entry^61–63^. At 24 hours p.i., the appearance of small MPXV foci was observed, visualized as groups of adjacent, virus-containing cells (Fig. 2b and c). In the following hours p.i., infection foci were observed increasing in size, while new virus-harboring cell clusters appeared (Fig. 2b and c). While infected NPCs appeared in compact rosette-shaped groups, in forebrain neurons fluorescence signal was detected additionally lying within neurites protruding from MXPV foci (Fig. 2b and c, Supplementary Movies 1 and 2).

Neural cell susceptibility to MPXV infection was further confirmed via transmission electron microscopy of organoids collected 10 days p.i. Pleomorphic, intracellular mature virions, characterized by a cuboid or ovoid structure with a central biconcave core were clearly visible within cells of the analyzed tissue (Fig. 2d’), and represented the majority of observed viral particles. Intracellular mature virions were visualized both in proximity to viral factories, and approaching the cells’ periphery, including regions of cell-to-cell contact. In addition, perinuclear viral factories, containing viral particles at different stages of the viral life cycle, were observed on multiple occasions (Fig. 2d’’). Viral crescent membranes at several stages of maturation were observed, along with immature viral particles, characterized by a smooth membrane encircling material of variable density (Fig. 2d’’), previously suggested to indicate different stages of virion organization^64^. Condensed cytoskeletal components were frequently documented in proximity of viral factories (Fig. 2d’’). Infected cells were commonly observed in clusters, and displayed features described to appear at advanced stages of viral replication^64^, including immature viral particles not only in nuclear proximity but also approaching host cell membrane, sustained nuclear morphology, and cytoskeletal condensations. In addition, we identified cells displaying evident signs of MPXV cytopathic effect, with nuclear degeneration and membrane rupture, leading to virus release (Fig. 2e). Extracellular virions without any evident association with cell structures were occasionally observed within hNO tissue. Further stages of viral particle maturation, including intracellular enveloped virions and cell-associated enveloped virions, were not identified.

### Monkeypox virus spreads from cell-to-cell in human neural cells

The vaccinia virus has been described exploiting several mechanisms to spread from cell-to-cell, including the formation of actin tails^65,66^ and tunneling nanotubes (TNTs)^67^. Driven by the previously observed presence of MPXV antigen within cell extensions of 2D cultured and organoid-enclosed neurons, we sought to further analyze the nature of the observed phenomenon. Consistent with previous observations, through transmission electron microscopy, we captured viral particles localizing not only to cell somata, but also to tube-like cellular extensions, rich in cytoskeletal components, reminiscent of neurites (Fig. 3a). In line with this observation, detailed immunofluorescence analysis of MPXV-challenged hNOs revealed infection not only limited to cell somata, but extending into thin cell filaments, frequently spanning between two cells within infection foci (Fig. 3b) and disseminated within the organoids’ cell meshwork. To elucidate the nature of the documented tubular structures, we performed co-localization analysis of viral antigen signal with F-actin and neuronal marker TUJ1, allowing discrimination between TNT-like structures and neurites^68^. Infected filaments of different length staining positive for both F-actin and TUJ1 were detected spanning between cells, confirming MPXV localization within neurites (Fig. 3c, yellow arrowheads). In addition, viral antigen accumulation within F-actin positive, but TUJ1 negative cell-connecting filaments, was occasionally observed (Fig. 3c, blue arrowheads), suggesting MPXV dissemination into, and possibly through, both neurites and TNTs. Intrigued by the observed viral distribution within cell-connecting neurites, we set out to further investigate the possibility of viral neuron-to-neuron transmission through dendrites and axons, a modality previously described for several neurotropic virus families, as reviewed by Taylor M.P. and Enquist L.W.^69^, but to our knowledge undocumented for *Poxviridae*. 2D cultured forebrain neurons were exposed to the MPXV-mNG reporter strain at an MOI of 0.01 TCID_50_/cell and imaged at regular intervals. Expression of mNG was observed to emerge in infected neurons within a few hours following infection, indicating ongoing viral DNA replication and protein translation. During this time, fluorescent signal was visualized within the cell somata and neurites of single cells scattered amidst the dense neuron population, in accordance with previously obtained results (Fig. 3d, Supplementary Fig. 4a and b, Supplementary Movies 2-4). As the infection progressed, mNG expression was frequently observed to subsequently appear in neighboring cells connected through signal-harboring filaments, often extending to several associated cells over time, substantiating neuron-to-neuron MPXV transmission (Fig. 3d, Supplementary Fig. 4a and b, Supplementary Movies 2-4).

**Figure 3:**
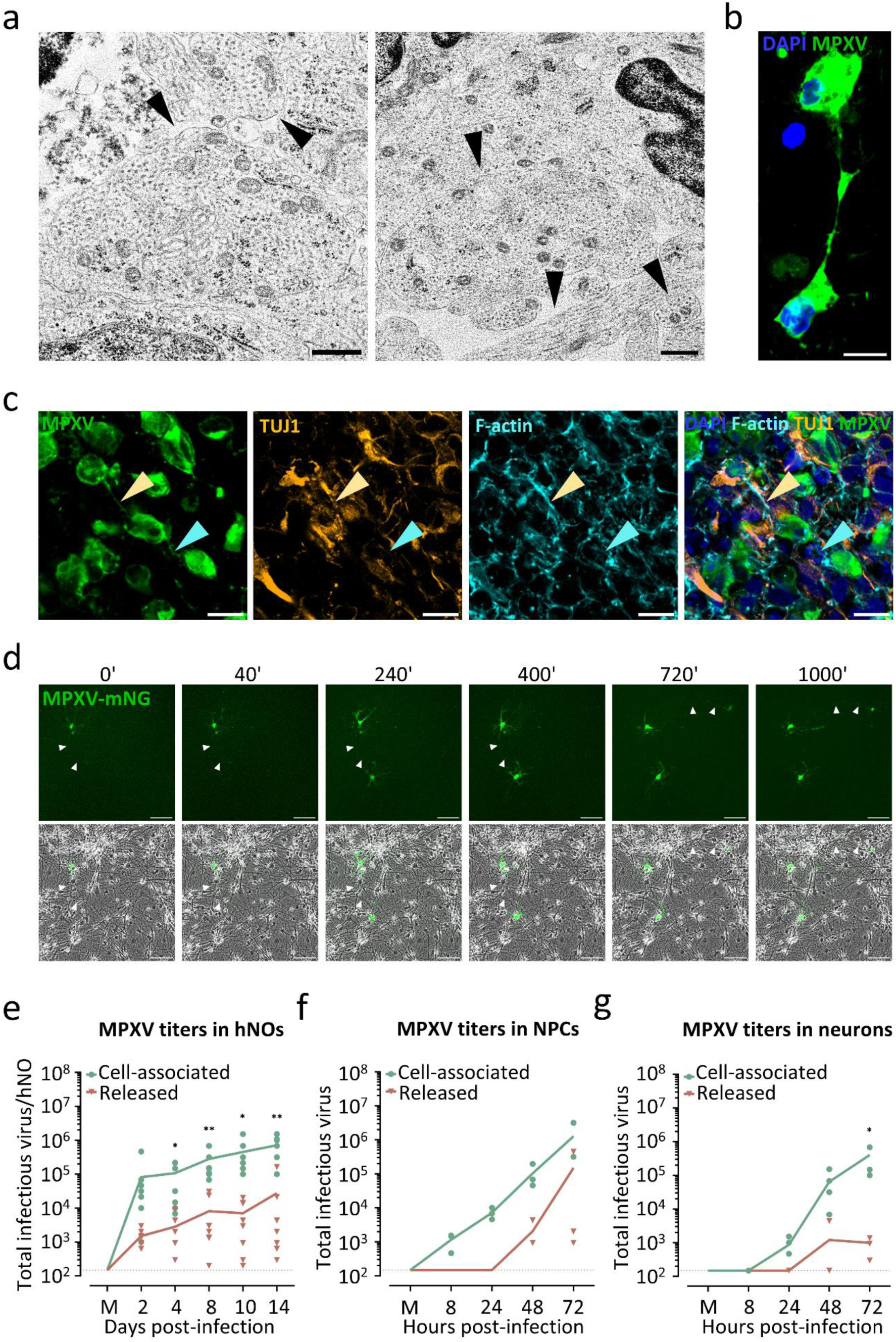
MPXV spreads through cell-to-cell contact. (a) Representative images acquired by transmission electron microscopy at 10 days p.i. of 95 days old hNOs exposed to a clade IIb lineage MPXV isolate at an MOI of 0.1 TCID_50_/cell. MPXV particles were observed localizing to neurites in infected hNO tissue. Black arrowheads indicate exemplary virion-harboring neurites. Scale bar, 0.5 µm. (b) Representative image of the detection by immunofluorescence of MPXV signal within neural filaments of interconnected, distinct cells. DAPI, blue; MPXV, green. Scale bar, 10 µm. (c) Representative micrographs of 95 days old hNOs infected with a clade IIb lineage MPXV isolate, showing the filaments connecting MPXV antigen positive cells and displaying diverse TUJ1 and F-actin patterns. Some filaments were observed showing expression of both markers (yellow arrowheads), indicative of their neuritic nature, others lacked TUJ1 signal, indicative of TNTs (blue arrowheads). DAPI, blue; MPXV, green; TUJ1, orange; F-actin, cyan. Scale bar, 10 µm. (d) Representative micrographs of 2D cultured forebrain neurons exposed to an mNG-expressing clade IIb lineage MPXV reporter strain, showing viral dissemination between interconnected neighboring cells. White arrowheads indicate mNG-harboring neurites. Images were taken at 40-minute intervals, time represented in minutes. Scale bar, 100 µm. (e) Cell-associated and released infectious MPXV 2 to 14 days p.i. of 70-95 days old hNOs infected with a clade IIb lineage isolate at a rate of 0.1 TCID_50_/cell. Each symbol represents an individual organoid (n = 7-8 independent organoid batches). Unpaired student’s t-test was applied to compare groups. *p < 0.05, **p < 0.01. (f) Cell-associated and released MPXV titers in 2D cultured NPCs challenged with a clade IIb lineage MPXV isolate at a rate of 0.01 TCID_50_/cell. Each symbol represents one NPC batch (n = 3 independent NPC batches). Unpaired student’s t-test was applied to compare groups. *p < 0.05, **p < 0.01. (g) Infectious MPXV released and cell-associated in 2D cultured neurons exposed to a clade IIb lineage MPXV isolate at a rate of 0.01 TCID_50_/cell. Each symbol represents one neuron batch (n = 4 independent neuron batches). Unpaired student’s t-test was applied to compare groups. *p < 0.05, **p < 0.01.

Having confirmed neurite-mediated virus transmission and considering the previously described ability of orthopoxviruses to spread via cell-to-cell contact rather than through extracellular-dissemination^65–67^, we set out to evaluate the extent of cell-associated transmission in both 3D and 2D environments. To discriminate between released and cell-associated infectious virus, we performed titrations of supernatants and homogenized hNOs, NPCs, forebrain neurons and Vero E6 cells. Within H1-derived organoids, cell-associated viral titers were observed to exceed released infectious virus at all measured time points, with differences spanning over several orders of magnitude (Fig. 3e). Significantly higher cell-associated than released viral concentrations were detected across all analyzed batches from day 4 p.i. on (Fig. 3e). In addition, cell-associated virus consistently contributed more than 85% of total infectious virus, with percentages fluctuating between 87% and 98% and steadily increasing over time, reaching the maximum at day 14 p.i. Comparable noteworthy, though not significant, differences were detected when IMR90-challenged organoids were analyzed (Supplementary Fig. 4c). Similar trends were documented for all 2D cultured cells, with intracellular viral titers constantly spanning above released amounts of infectious virus. In neural cell types, growth curves of extracellular and cell-associated titers were observed converging towards the end of infection in NPCs, while infectious viral titer differences between the two analyzed compartments were documented increasing over time in neuronal cultures (Fig. 3f and g). Consistently, the divergence between released and cell-associated titers was observed to be important though not significant in NPCs at all analyzed time points, whereas significance was confirmed at 72 hours p.i. in neuronal cultures (Fig. 3f and g). While neuron cultures showed a delay in MPXV replication within the intracellular compartment, after 8 hours p.i. cell-associated virus dynamics between NPCs and neuron cultures were deemed comparable (Fig. 3f and g). In contrast, released infectious virus titers showed noticeable differences between the two analyzed cell types. While extracellular MPXV was observed only slightly increasing between 24 and 48 hours p.i. in neuron cultures and plateauing afterwards, starting at 24 hours p.i. virus amounts released by NPCs displayed a constant increase, further corroborating a selective preference of MPXV for neurite-associated transport (Fig. 3f and g). In accordance with light-microscopy observations, highly susceptible Vero E6 cells displayed rapidly increasing viral titers in the intracellular compartment from 24 till 72 hours p.i., after which no further increase was recorded (Supplementary Fig. 4d). While a delay of 24 hours was observed in released amounts of infectious virus, comparable dynamics to the ones previously seen in NPCs were subsequently recorded (Supplementary Fig. 4d). Despite the strong increase of released infectious virus, however, significant differences between the two virus-harboring compartments were found both at 72 hours p.i. and 96 hours p.i. (Supplementary Fig. 4d).

### Monkeypox virus infection leads to neurite injury

Further examination of MPXV-infected hNO cultures revealed viral antigen signal accumulating at regular intervals within elliptical swellings along infected filaments (Fig. 4a). Notably, these structures were observed in 100% of analyzed independent batches of H1- and IMR90-derived hNOs. Analogous shape modulations, termed neuritic (i.e., axonal or dendritic) beads, have been postulated to indicate degenerative processes and reported to accompany a variety of neuropathologies including Alzheimer’s disease^70–72^, ischemia^73^, and traumatic injuries^74^. Beaded filaments were documented spanning for remarkable distances throughout the hNO tissue, both within and outside of infection foci, and swellings were observed to be present in both symmetric, lemon-shaped and clam-shaped, asymmetric morphology, indicative of early and advanced stages of beading^75^. Detailed analysis showed frequent co-localization of viral antigen accumulations with neuronal marker TUJ1, both within swellings and affected filaments, suggesting gathering of viral proteins along dendrites and axons (Fig. 4b). The neuritic nature of beaded filaments was further corroborated by staining with alternative neuron markers, including microtubule-associated protein 2 (MAP2)^76^ and transcription factor neuronal nuclei (NeuN)^77^ (Supplementary Fig. 5a and b). Aiming to document the formation and injurious nature of MPXV-induced neuritic beading, 2D cultured forebrain neurons were exposed to MPXV-mNG and recorded through live-cell imaging at regular intervals. Bead formation was confirmed to represent a common phenomenon in MPXV-harboring neurons frequently recorded in the days following the first appearance of fluorescent signal (Fig. 4c, Supplementary Fig. 5c, Supplementary Movies 4-6). Intriguingly, reporter expression was observed to abruptly disappear from neurites and subsequently somata shortly after bead appearance, with fluorescent signal accumulating within the previously formed beads (Fig. 4c, Supplementary Fig. 5c, Supplementary Movies 4-6). Somata of affected neurons were furthermore observed to undergo major structural changes around the time of bead appearance, including bleb formation (Fig. 4c, Supplementary Fig. 5c-e, Supplementary Movies 4-6), cell shrinkage (Supplementary Fig. 5d), and cell rounding and detachment (Supplementary Fig. 5e), accompanied by mNG signal fading, indicative of cell degeneration. Previously, enrichment of the pro-apoptotic protein cleaved caspase-3 (CC3)^78,79^, had been reported in degenerating neurons^80,81^. We thereby set out to analyze the distribution of CC3 within beaded neurons in infected hNOs. In line with previous reports, we observed occasional CC3 accumulation in neuritic beads as well as within associated cells (Supplementary Fig. 5f).

**Figure 4:**
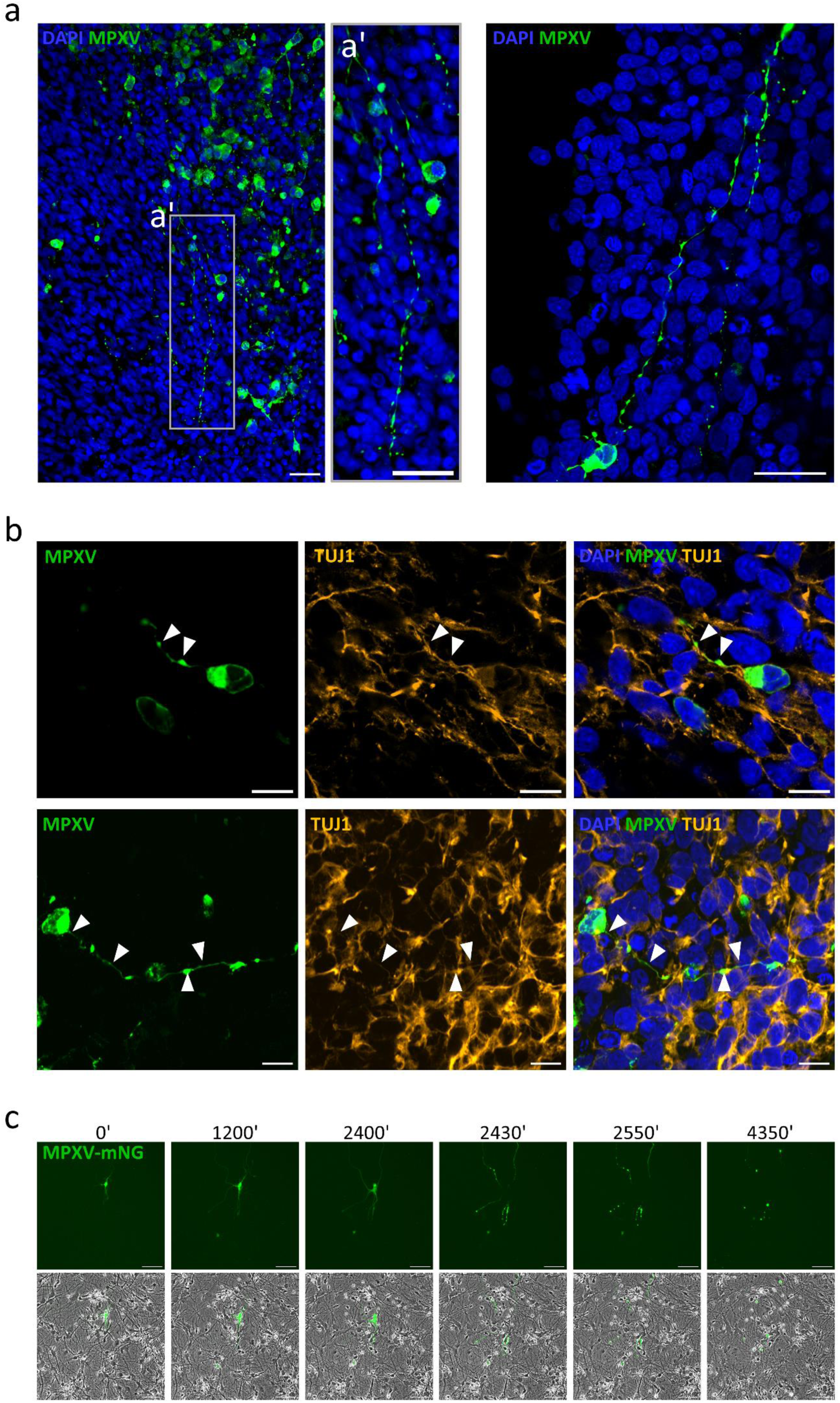
MPXV causes neuritic beading and neuron death. (a-b) Immunofluorescence analysis 8-14 days p.i. of 70 days old hNOs exposed to a clade IIb lineage MPXV isolate at a rate of 0.1 TCID_50_/cell. (a) Representative micrographs showing viral antigen accumulating within regularly interspaced swellings (beads) on filaments, spanning over long distances within the tissue. DAPI, blue; MPXV, green. Scale bar, 25 µm. (b) Representative micrographs of beaded filaments harboring monkeypox virions co-localizing with the neuronal marker TUJ1 in both filaments and beads. White arrowheads indicate regions displaying marker co-localization. DAPI, blue; MPXV, green; TUJ1, orange. Scale bar, 10 µm. (c) Representative micrographs of 2D neurons exposed to an mNG-expressing clade IIb lineage MPXV reporter strain displaying neuritic beading on filaments of virus-harboring cells in concomitance with sudden mNG signal loss in cell soma. mNG signal is subsequently observed to progressively diminish while beads still harbor detectable reporter protein signal. Images were taken at 30-minute intervals, time represented in minutes. Scale bar, 100 µm.

### Monkeypox virus disrupts cellular homeostasis and leads to synaptic dysregulation

Reports from the 2022 MPXV outbreak indicate detection of several inflammatory cytokines and chemokines in mpox patients^82,83^. Aiming to characterize the innate immune response elicited by MPXV-challenge in hNOs, we analyzed the regulation of four selected inflammatory mediators, including interleukin-6 (IL-6), interleukin-8 (IL-8/CXCL8) and interferon gamma-induced protein 10 (IP-10/CXCL10), previously found to be significantly increased in mpox patients^82,83^, as well as interferon-β (IFN-β), a central effector cytokine of the host antiviral immune response^84–86^. In line with previous reports, quantitative PCR (qPCR) results revealed upregulation of all analyzed cytokines, at all evaluated time points (Fig. 5a). A rapid, significant increase in IL-6, IL-8 and IP-10 was observed at 4 days p.i., while IFN-β showed a noticeable, though not significant upregulation, indicating early induction of inflammatory responses and establishment of antiviral state in our cultures (Fig. 5a). Elevated mRNA levels were maintained through day 10 p.i. for all analyzed cytokines, though inter-organoid variations were observed (Fig. 5a).

**Figure 5:**
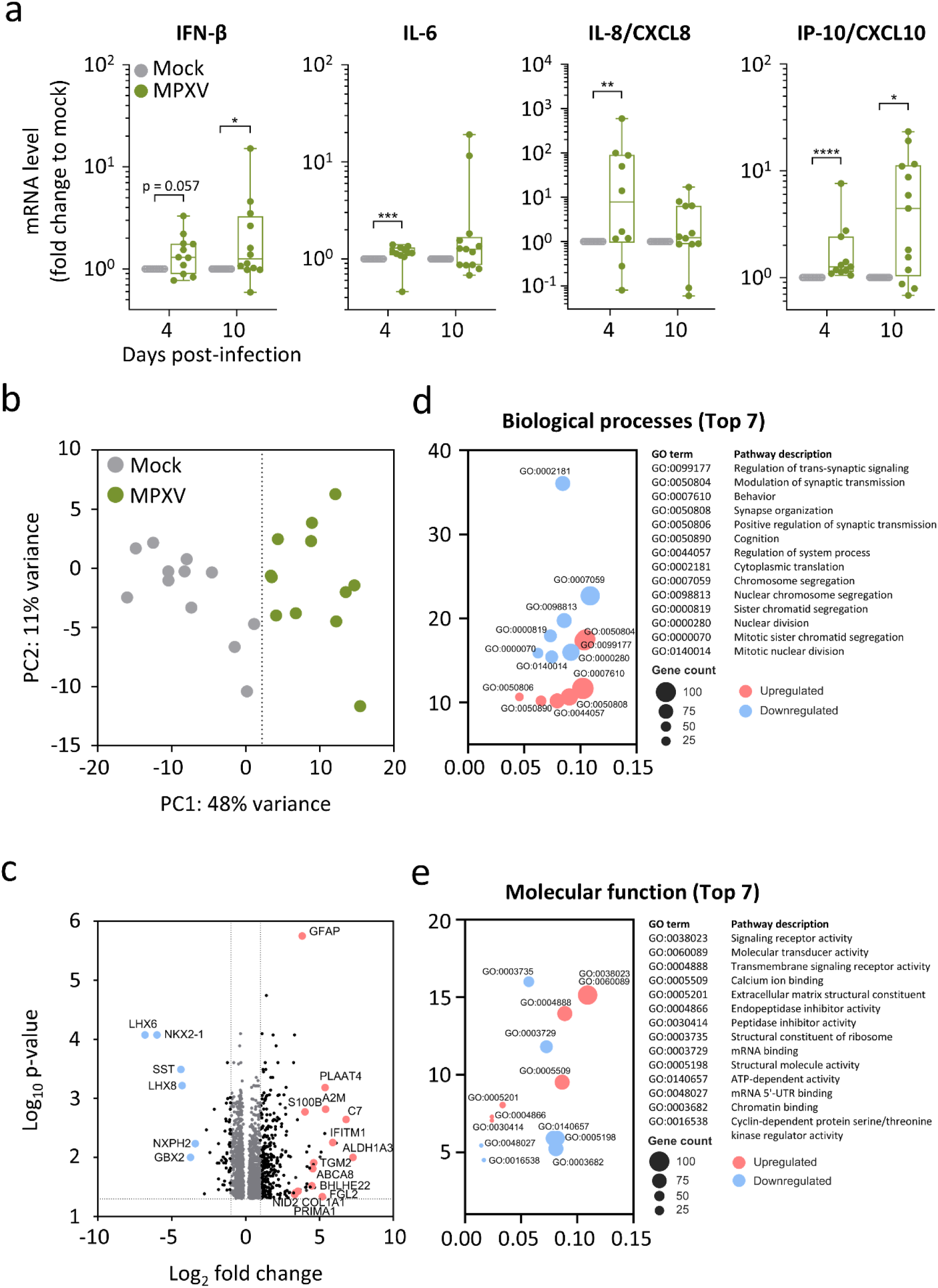
Transcriptional profile alteration in hNOs during MPXV infection. (a) Fold change of transcriptional expression levels of selected proinflammatory cytokine and chemokine genes including IFN-β, IL-6, IL-8/CXCL8 and IP-10/CXCL10 in 74-93 days old hNOs exposed to an MPXV clade IIb lineage isolate at a rate of 0.1 TCID_50_/cell. Boxplots indicate the median value (centerline) and interquartile ranges (box edges), with whiskers extending to the lowest and the highest values. Each symbol represents an individual organoid (n = 3 independent organoid batches). Mann-Whitney U test was applied to compare groups. *p < 0.05, **p < 0.01, ***p < 0.001, ****p < 0.0001. (b-e) Transcriptomic analysis of MPXV-challenged versus mock-treated hNOs infected with a MPXV clade IIb lineage isolate at a rate of 0.1 TCID_50_/cell at 74-93 days of development (n = 3 independent organoid batches). hNOs were collected for RNA isolation at 4 days p.i. (b) PCA showing separate clustering of mock and MPXV-exposed organoids. (c) Volcano plot representing DEGs between mock-treated and MPXV-challenged hNOs; p_adj_ < 0.05. (d) GO analysis results displaying top 7 up- and downregulated biological processes in mock-treated versus MPXV-infected hNOs; p_adj_ < 0.05. (e) GO analysis results displaying top 7 up- and downregulated molecular functions in mock-treated and MPXV-exposed hNOs; p_adj_ < 0.05.

Seeking to delve deeper into MPXV-induced host responses in human neural tissue, the transcriptome of mock-treated and MPXV-infected hNOs at day 4 p.i. was analyzed through bulk RNA sequencing (Fig. 5b-e). Principal component analysis (PCA) of differentially expressed genes (DEGs) revealed separate clustering of mock-treated and MPXV-exposed hNOs (Fig. 5b). Differential expression analysis showed significant downregulation of 871 and induction of 921 transcripts following MPXV infection (Fig. 5c). Surprisingly, transcripts indicative of ongoing antiviral and inflammatory processes, as the interferon-induced transmembrane protein 1 (IFITM1)^87,88^ and fibrinogen-like protein-2 (FGL-2)^89^, represented only a minority of DEGs (Fig. 5c). In contrast, numerous genes associated with neurodegenerative process modulation, as the alpha-2 macroglobulin (A2M), associated with neuronal injury in Alzheimer’s disease^90^, and S100 calcium-binding protein (S100B), reported to be upregulated in a variety of brain pathologies^91^, were identified amongst the most highly upregulated DEGs (Fig. 5c). In line with our observations, gene ontology (GO) analysis for biological processes revealed strongest enrichment of pathways related to neuronal reorganization, including “regulation of trans-synaptic signaling” and “synapse organization” (Fig. 5d). Furthermore, GO terms pertinent to behavior and cognition were identified as highly enriched under MPXV influence (Fig. 5d). In contrast, terms related to cell division and housekeeping activities, as “cytoplasmic translation” and “chromosome segregation”, were observed dominating downregulated biological processes, indicating inhibition of cell division and disruption of cell homeostasis upon MPXV-infection in hNOs (Fig. 5d). GO molecular function analysis furthermore revealed upregulation of cell signaling processes along with terms hinting towards changes in tissue organization, as “signaling receptor activity” and “extracellular matrix structural constituent” (Fig. 5e). Consistent with previous observations, terms associated with basic cellular functions, as “structural constituent of ribosome” and “chromatin binding” were amongst the most importantly attenuated molecular functions (Fig. 5e). Overall, while a steady upregulation of inflammatory mediators was observed through targeted mRNA analysis, our findings reveal a transcriptional landscape dominated by tissue reorganization in hNOs facing MPXV-infection.

### Tecovirimat treatment limits monkeypox virus spread in human neural organoids

Use of repurposed smallpox antiviral drugs as tecovirimat and cidofovir currently represents one possible line of treatment for patients suffering from severe mpox courses^92^. Due to scarcity of clinical evidence however, to date, their efficacy remains inconclusive^92,93^. To evaluate whether tecovirimat could represent a possible treatment approach for patients affected by mpox-encephalitis, we set out to analyze the antiviral activity of tecovirimat in hNOs. Organoids were subjected to MPXV as previously described, and antiviral treatment was initiated 2 hours after virus exposition. To evaluate optimal treatment dose, one batch of hNOs was exposed to medium supplemented with increasing concentrations of tecovirimat, starting at 0.1 μM till 5 μM, with concentrations of 1 μM and 5 μM representing clinically significant plasma concentrations^94–96^. A dose-dependent reduction of cell-associated infectious viral loads was observed at all analyzed time points, with the lowest fold-change observed 14 days p.i. in hNOs exposed to 0.1 μM tecovirimat, and the highest decrease at 8 days p.i. when 5 μM were applied (Supplementary Fig. 6a). We opted to proceed further investigations using the intermediate and clinically relevant dose of 1 μM. Three independent batches of hNOs were challenged with MPXV and treated with the previously established protocol. No impact of tecovirimat exposure on hNO cultures was observed by light microscopy analysis of organoid outer morphology after 14 days of treatment, indicating no evident toxic effect of drug exposure on neural tissue (Fig. 6a). Furthermore, inner morphology analysis of sectioned and immunofluorescence-stained organoids, processed as previously described, confirmed no disruption of tecovirimat treatment on hNO cell organization (Fig. 6b). The limiting effect of tecovirimat application on MPXV replication was confirmed by confocal microscopy. While increasing MPXV-antigen accumulation was previously detected over the course of infection in MPXV-challenged organoids, tecovirimat-exposed organoids displayed only sporadic appearance of virus-harboring cells and infection foci at all analyzed time points (Fig. 6b). Despite the important decrease of antigen-carrying cells, however, tecovirimat treatment did not prevent formation of neuritic beads in hNOs (Supplementary Fig. 6b). Quantification of infectious virus levels substantiated visual findings, confirming a strong reduction of both released and cell-associated viral titers at all analyzed time points (Fig. 6c and d). While no released infectious virus was detectable through titration in tecovirimat-treated cultures, cell-associated viral titers showed decreased total infectious viral loads at all timepoints (Fig. 6d). Cell-associated infectious virus was observed decreasing with length of treatment, showing only a 2.3-fold decrease on the second day of infection, but a 19.5-fold decrease by day 14 p.i. (Fig. 6d). To exclude possible inhibition of plaque-forming capacity of MPXV on Vero E6 cells by residual tecovirimat in the context of titration, viral load reduction was furthermore confirmed through qPCR analysis (Fig. 6e and f). While released MPXV DNA was still detectable through qPCR analysis, its level was importantly decreased at all time points (Fig. 6e). Similarly, qPCR of cell-associated MPXV DNA contents confirmed previous observations, with viral DNA loads in tecovirimat-treated organoids spanning several orders of magnitude below unexposed cultures (Fig. 6f). Taken together, our results indicate that relevant reduction of MPXV infection in neural tissue can be achieved through tecovirimat treatment in both extracellular and intracellular compartments, mitigating neuronal injury, though not completely rescuing it.

**Figure 6:**
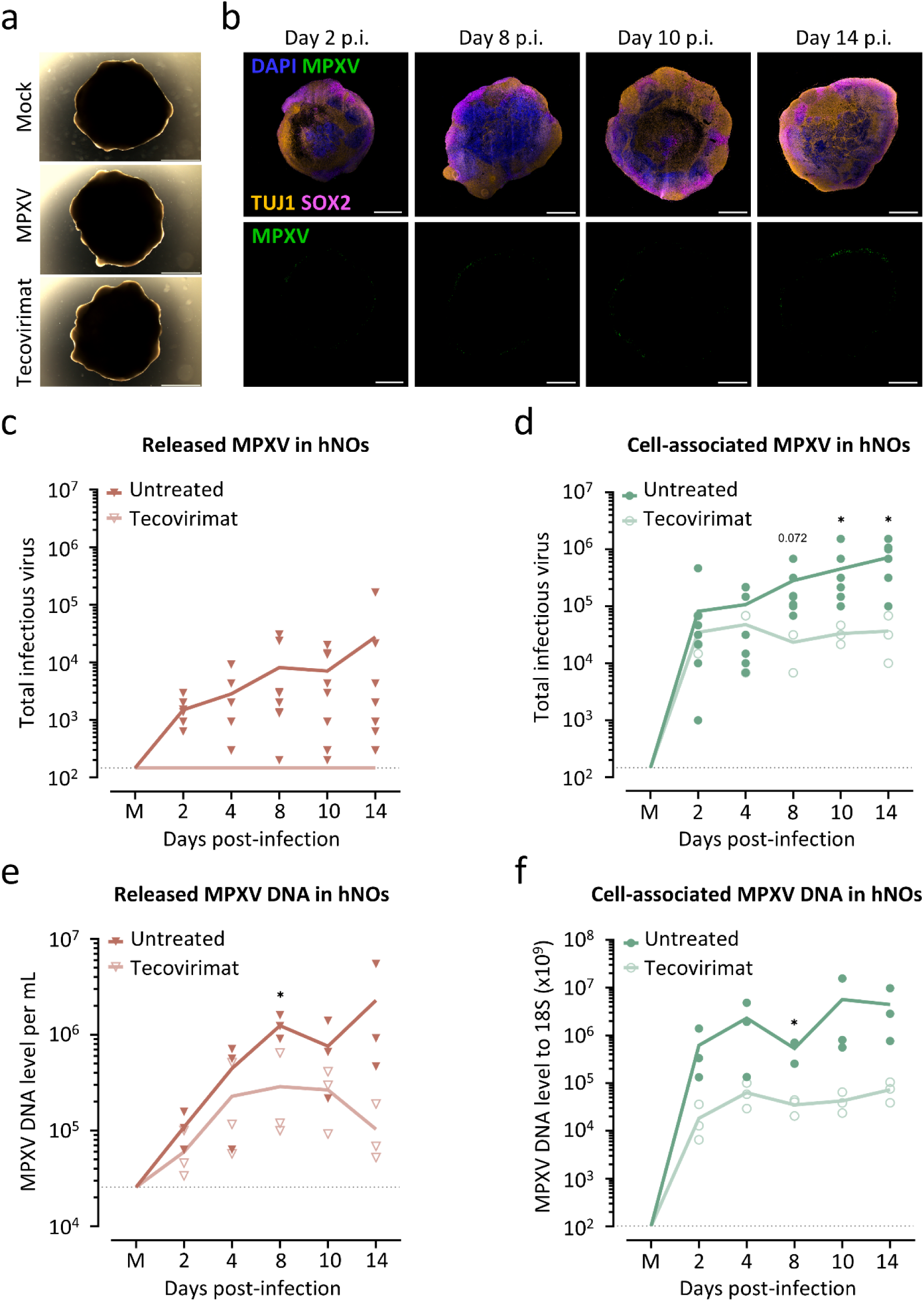
Tecovirimat treatment limits MPXV spread in hNOs. (a) Representative micrographs of mock-treated (top) and MPXV-exposed hNOs cultured in regular (center) or tecovirimat complemented medium (bottom) at 14 days p.i. Organoids were exposed to a clade IIb lineage MPXV isolate at a rate of 0.1 TCID_50_/cell at 84 days of development. Tecovirimat was added to the medium 2 hours after initial MPXV-exposure, to a final concentration of 1 µM. Scale bar, 2000 μm. (b) Representative sections of tecovirimat-treated hNOs sampled at selected time points p.i. showing unaltered tissue organization and limited virus antigen presence. Infected organoids were exposed to a clade IIb lineage MPXV isolate at a rate of 0.1 TCID_50_/cell at 84 days of development. DAPI, blue; MPXV, green; TUJ1, orange; SOX2, pink. Scale bar, 1000 μm. (c-d) Released (c) and cell-associated (d) infectious MPXV 2 to 14 days p.i. of 84-85 days old hNOs infected with a clade IIb lineage isolate at a rate of 0.1 TCID_50_/cell and cultured in regular or tecovirimat-supplemented medium. Each symbol represents an individual organoid (n = 3 independent organoid batches). Unpaired student’s t-test was applied to compare groups. *p < 0.05, **p < 0.01. (e-f) Released (e) and cell-associated (f) MPXV loads detected through qPCR analysis in samples taken 2 to 14 days p.i. 84-85 days old hNOs were exposed to a clade IIb lineage MPXV isolate at a rate of 0.1 TCID_50_/cell and cultured in regular or tecovirimat-supplemented medium. Each symbol represents an individual organoid (n = 3 independent organoid batches). Unpaired student’s t-test was applied to compare groups. *p < 0.05, **p < 0.01

## DISCUSSION

In this study, we explored the neurobiology of MPXV in the human host using hNOs as an advanced model of the human brain. Our data provides evidence that several neural cell types are susceptible to MPXV infection, and the resulting replication leads to virus spread throughout the neural tissue. Moreover, evaluation of infectious virus titers in intra- and extracellular compartments indicates a spread of MPXV via cell-to-cell contact in hNOs, 2D NPCs and neurons. Through live-cell imaging analysis, we show repurposing of axons and dendrites in 2D forebrain neuron cultures as highways for MPXV spread. The identification of neuronal injury hallmarks and neurodegenerative signs demonstrates the neurovirulence potential of MPXV in humans. We furthermore show that MPXV infection leads to significant changes in the transcriptome of hNO tissue, driven towards synaptic reorganization and confirming upregulation of neurodegeneration-associated genes, while presenting a significant suppression of housekeeping processes. In line with clinical data, we confirm limitation of MPXV spread upon organoid exposure to tecovirimat, hinting to possible treatment options for mpox-affected encephalitis patients.

Despite MPXV infection cases being reported since 1970, studies investigating pathomechanisms driving mpox disease are sparse. More so, current understanding of processes leading to rare, but often severe complications in mpox-affected individuals, shows prominent gaps. We used hNOs to model MPXV infection in the human host, as we confirmed that prolonged culture allows establishment of relevant cell diversity, within a highly complex environment. We show that hNOs are susceptible to and allow active replication of MPXV, as demonstrated by transmission electron microscopy, immunofluorescence analysis and increasing cell-associated and released infectious virus. Transmission electron microscopy of hNOs revealed presence of intracellular viral particles at several stages of viral life cycle, including crescent membranes, immature viral particles, intracellular mature viral particles, and extracellular viral particles. These findings show the establishment of MPXV replication machinery in human neural tissue is possible and frequently occurs within infected hNO cells. As previously described in Vero cells^64^, we observed abundance of intracellular mature virions over all other viral morphotypes, and documented their presence at different subcellular locations, including in close proximity to viral factories, possibly indicating recently formed virions, and adjacent to cell membranes. Furthermore, we confirmed the recurrent presence of bundled cytoskeletal elements, which we frequently observed located in proximity to viral factories. Cytoskeletal rearrangements have been recently reported in MPXV-infected Vero cells^64^. In addition, cytoskeleton involvement has been suggested in processes mediating viral distribution within infected cells^59,65,66,97,98^, as well as during cell-to-cell transmission processes, as actin tail formation^65,66,98^. We thus hypothesize that cytoskeletal condensations observed within hNO cells might be involved in MPXV transport within the cells’ cytoplasm.

In line with the establishment of a functional viral replication machinery within hNO cells, immunofluorescence analysis showed a steady viral antigen increase in the course of infection, reaching a peak around day 10 p.i. Viral antigen was initially visualized on the organoids’ surface, occasionally reaching deeper into the tissue at advanced stages of infection. In addition, viral particles were often detected localizing to distinct cell foci or streams, suggesting possible cell-to-cell transmission, a prominent route exploited by orthopoxviruses, as vaccinia virus^65–67^. While no actin tails were visualized by transmission electron microscopy, intracellular mature viral particles were observed within neurite sections in the processed tissue. In addition, immunofluorescence analysis showed viral antigen localizing not only to cell somata, but also to filaments of variable length, connecting distinct cells and dispersed within the organoid tissue. Characterization of cell processes showed the presence of both TUJ1-positive neurites and F-actin-positive, TUJ1-negative TNT-like structures harboring viral particles within hNOs, suggesting that cell-to-cell transmission might occur through multiple routes. Fluorescence microscopy and live-cell imaging of 2D neurons exposed to a MPXV clade IIb mNG-reporter construct further confirmed our findings, showing mNG signal in neurites early after infection onset, followed by the infection of neighboring, filament-connected cells. Comparison of cell-associated and released infectious virus in 2D cultured cells and 3D tissue further strengthened this assumption. Indeed, intracellular viral loads were found to exceed the amount of released virus in all models. However, while both 2D NPCs and forebrain neurons were found accumulating infectious viral particles within intracellular compartments, significant differences were recorded for neuron cultures only, further consolidating the relevance of axons and dendrites as an additional route for MPXV cell-to-cell transmission. In accordance, through live-cell imaging, we recorded infection propagation between interconnected neurons in 2D cultures. Together, our findings suggest that, as previously documented for the closely related vaccinia virus, MPXV preferentially spreads directly from cell-to-cell and can exploit different mechanisms to spread within human neural tissue. Furthermore, while our observations confirm the possible transmission of MPXV through TNT-like structures, as earlier documented for vaccinia virus^67^, we predominantly detected localization of viral antigen within TUJ1-positive neurites in hNOs and found a significant enrichment of cell-associated infectious MPXV in forebrain neuron cultures. These observations suggest a previously undescribed mode of viral transmission for MPXV in the human brain, in which viral particles can be transported from one cell to another through axons and dendrites. Documented cytoskeletal alterations in proximity to viral factories could hint at the mechanism of transportation of the virus, which could exploit the cells’ fast axonal transport machinery to move through the cytosol and reach distant locations, a mechanism observed to be exploited by viruses of several families, most prominently *Herpesviridae*, *Flaviviridae*, *Rhabdoviridae* and *Picornaviridae* as reviewed by Taylor M.P. and Enquist L.W.^69^, allowing viral dissemination while evading the host’s immune system. Various mechanisms driving cytoskeletal manipulation by the closely related vaccinia virus have been previously demonstrated, encompassing both actin^65,66,99–101^ and microtubule-based^59,102,103^ pathways to increase pathogen motility. In agreement with these observations, cytoskeletal reorganization was recently shown in primary human fibroblasts infected with a clade II MPXV isolate^104^, further corroborating our hypothesis.

In addition, our findings reveal the injurious consequences of viral spread within neurites. Viral antigen-containing filaments within hNOs were often observed showing sequential swellings, termed neuritic beads. Consistently, we detected bead formation in neurites of 2D cultured neurons. Bead appearance was additionally observed to precede or coincide with striking changes in cell morphology, including membrane blebbing and cell shrinkage or rounding and detachment, indicative of neuronal death. Analogous bead formation has been reported to accompany diverse pathological states, including neurodegenerative disorders such as Alzheimer’s disease^70–72^, ischemia^73^, traumatic injuries^74^ and neurotropic virus infections^105–107^. Beads are thought to originate from changes in membrane tension, and organelles, cytoskeletal components as well as pathology-associated proteins have been observed to sporadically pause or accumulate within them^75,108^. Furthermore, beads have been observed to appear in axons of trigeminal neurons upon pseudorabies virus cell attachment^109^. These swellings have been suggested to represent sites of virus egress^109^ and could possibly be accompanied by a deleterious interference with regular axonal functions^110^. It seems reasonable to hypothesize that MPXV hijacks the cells’ transport machinery, causing cytoskeletal rearrangement, and exploits it to move through axons, leading to the generation of neuritic beads that might represent sites of viral egress, finally causing neuronal injury by hampering cell functions. Our findings furthermore indicate that sudden death of MPXV-infected neurons can be triggered merely by viral presence. We additionally report occasional but inconsistent accumulation of CC3 within neuritic beads and parental neurons. The extent to which apoptosis contributes to the observed phenomenon and whether other mechanisms are involved requires further clarification.

Recently, Chailangkarn and colleagues^55^, found iPSC-derived NPCs and astrocytes to be susceptible to MPXV infection in a 2D culture environment. This was demonstrated by changes in cell morphology and viability, as well as increased viral loads detected in supernatant by qPCR over time^55^. In contrast, 14 and 45-day old neurons did not show productive MPXV infection^55^. In an unrefereed preprint, human pluripotent stem cell-derived cortical neurons were however shown to be permissive to MPXV infection by Bauer and colleagues^111^, even if at a lower degree compared to astrocytes and microglia. In line with both reports, we confirmed susceptibility of both NPCs and astrocytes to MPXV infection and found TUJ1-expressing neurons to be permissive to MPXV infection as well. We further corroborated our findings through 2D analysis of NPC and forebrain neuron cultures, with infectious virus quantification showing an increased preference of MPXV cell-to-cell spread in neurons. We propose that the prevalence of cell-associated neuronal spread of MPXV through axonal transport provides a rationale for the previously reported low or undetected viral replication within human neuron cultures, suggested to resist viral infection to a certain degree, based on the lack of a strong increase in detectable virus in culture supernatant. Further investigation of the exact mechanisms driving the observed phenomena could prove of utmost importance in the development and identification of novel treatment options for patients affected by mpox-encephalitis.

An intricate interplay of host-pathogen interactions modulates the course of viral infections, including the neuroinvasive potential of viruses and pathogenic consequences of CNS infection. Several studies have reported transcriptional changes induced by MPXV clade I infection in a diversity of models, including rhesus macaque kidney epithelial cells (MK2)^63^, HeLa cells^112,113^, primary human fibroblasts and macrophages^113^. Furthermore, MPXV clade IIb lineage-induced alterations were analyzed and reported in human skin organoids^95^, human kidney organoids^96^ as well as iPSC-derived neural cell types^111^. Results vary greatly between reports, suggesting cell-type specific responses to MPXV challenge. We investigated the consequences of MPXV neuroinvasion through hNO transcriptome analysis and targeted qPCR. In our study, significant induction of inflammatory cytokines and chemokines was confirmed through sensitive qPCR analysis, albeit a scarcity of innate-immunity related DEGs was noticed in the RNA sequencing data. MPXV and related members of the *Poxviridae* family present a highly complex genome, with several encoded proteins postulated to exert immunomodulatory functions, allowing immune response silencing and evasion^114,115^. In line with this, primary human monocyte-derived macrophages and primary human dermal fibroblasts have been reported to present a striking lack of response in spite of dramatic cytopathic changes when exposed to infectious clade I MPXV, due to selective viral host-response inhibition^113^. Our observations align with a sustained MPXV-driven suppression of host antiviral defense mechanisms. Interestingly, pathway analysis revealed significant upregulation of signaling-related terms, especially those associated with synaptic functions. Differential gene expression of synapse-associated cellular components had been previously recorded in MPXV-exposed HeLa cells^116^. Similar observations have been furthermore reported during West Nile virus infection in rhesus macaques, in which histological changes correlated with significant alteration of synaptic gene regulation^117^. Direct virus-mediated dysregulation of synaptic homeostasis as well as compensatory host mechanisms initiated upon MPXV-induced neuronal injury could underlay the observed effects of infection on synapse function. Alternatively, neuritic beads induced by α-herpesvirus infection have been reported to express the synapse marker synaptophysin and possibly represent functioning synapses^109,118^. MPXV-mediated formation of axonal and dendritic beads with synapse-like activity could hence account for the upregulation of synapse signaling-related terms, although this hypothesis appears less probable considering the rapid succession of events following bead induction leading to neuron degeneration. In our analysis, perturbation of neural cell homeostasis was further demonstrated by significant downregulation of genes related to basic cellular housekeeping functions, most prominently cell division. Similarly, modulation of cell cycle progression by MPXV clade I infection has been previously reported in challenged MK2 cells and has been suggested to represent an antiviral host strategy common to several viral pathologies^63^. Alternatively, host cell cycle arrest in favor of viral replication has been suggested for other poxvirus infections^104,119,120^. Overall, transcriptome analysis of MPXV-exposed hNOs reveals profound perturbation of neural tissue, predicted to potentially deeply impact brain function.

Current treatment options for patients affected by severe courses of mpox disease, included those affected by mpox-encephalitis, include supportive care, immunomodulatory therapies and antiviral drug administration, as tecovirimat or cidofovir^92^. While early treatment initiation was found to potentially reduce virus shedding duration^93^, evidence supporting efficacy of tecovirimat treatment in the 2022 outbreak is limited. Furthermore, preliminary data suggests that tecovirimat did not reduce the duration of mpox symptoms in patients infected with clade I MPXV^121^. However, tecovirimat has been shown to permeate the blood-brain barrier in several animal models^122,123^, thereby representing a possible line of treatment for mpox-encephalitis affected patients. We confirm a dose-dependent reduction of viable MPXV in tecovirimat-treated organoids and report that a prolonged drug exposition can significantly reduce infectious MPXV, while exerting no obvious toxic effects on hNO tissue. In line with the drug’s mode of action, known to prevent MPXV egress but not formation of intracellular mature virions^124^, however, neuritic beading was not ablated by tecovirimat treatment. These findings confirm that neurite-mediated transport of intracellular infectious MPXV can sustain virus spread in absence of extracellular enveloped viral particle production and show that released and cell-associated virus dissemination synergically contribute to efficient virus infection. Our findings furthermore indicate that, in spite of tecovirimat tampered virus propagation, injurious effects of sustained neuron-to-neuron transmission persist.

Taken together, we show that human neural tissue, modelled in a complex 3D environment, is susceptible to infection with a contemporary clade IIb lineage MPXV isolate. We show that viral replication factories are successfully established, resulting in a productive replication of MPXV within organoid cells. Furthermore, we find that viral antigen localizes not only to cell somata, but also to filaments of variable nature. We propose that MPXV preferentially spreads from cell-to-cell, exploiting not only previously described mechanisms, but also through axonal transport, as demonstrated through live-cell imaging visualization of virus propagation dynamics. We furthermore report neuritic bead formation in virus-harboring neurites and dendrites, previously documented to represent sites of virus egress and cell-to-cell transmission, as well as signs of neuronal injury. Notably, bead formation precedes virus-induced neuronal death, which is initiated through neurite degeneration. The transcriptional landscape of MPXV-infected neural cultures suggests repurposing of tissue to favor viral propagation, characterized by disrupted cell homeostasis and upregulation of transcripts associated with neurodegenerative processes and synaptic reorganization. Notably, tecovirimat treatment effectively limits viral spread but does not rescue the deleterious effects of neuron-to-neuron MPXV dissemination. Our findings identify a novel mechanism of MPXV spread in human neural tissue, shed light on possible factors contributing to mpox-encephalitis neuropathology and constitute the basis for further exploration of the neurobiology of orthopoxviruses.

## MATERIAL AND METHODS

### Ethics statement

In accordance with Articles 13 and 14 of the Federal Act on Research on Embryonic Stem Cells, and Article 20 of the Ordinance on Research on Embryonic Stem Cells, the work with human H1 ESC line was approved by the Cantonal Ethics Committee of Bern, Switzerland under the authorization number R-FP-S-2-0023-0000.

### Maintenance of human pluripotent stem cells

The human H1 ESC line (WiCell) and the human IMR90 iPSC line (WiCell) were cultured in feeder-free conditions and used to generate hNOs employed in this study. ESCs and iPSCs were seeded onto untreated culture flasks (Sigma-Aldrich) coated with Vitronectin XF (Stem Cell Technologies) and maintained in mTeSR Plus medium (Stem Cell Technologies). Cells were maintained according to the manufacturer’s recommendations. Cells were passaged every 6-7 days applying 0.5 mM EDTA (Thermo Fisher Scientific) in sterile Dulbecco’s Phosphate Buffered Saline (DPBS; Gibco) for 4 minutes at 37°C, 5% CO_2_, followed by mechanical dissociation to obtain aggregates encompassing approximately 5-8 cells. Regions with differentiated cells were scratched prior to cell detachment. Cells were regularly checked and found negative for mycoplasma contamination using the LookOut Mycoplasma qPCR Detection Kit (Merck).

### Generation and maintenance of human neural progenitor cells

Human NPCs were differentiated using STEMdiff Neural Induction Medium (Stem Cell Technologies) as per manufacturer’s instructions, with minor changes. Briefly, in the days preceding the generation or splitting of NPCs, cell-culture treated 6-well plates (Falcon) or 24-well μ-plates (ibidi) were coated. Poly-L-ornithin (PLO; Sigma-Aldrich) solution was diluted in DPBS to obtain a final concentration of 15 μg/mL. The solution was evenly distributed on the wells’ surface and left to incubate at room temperature or at 37°C for a minimum of 2 hours. Subsequently, cultureware was washed twice with DPBS and once with DMEM/F-12 (Gibco). A 10 μg/mL laminin (Sigma-Aldrich) solution in DMEM/F-12 was subsequently added to the PLO-coated wells and left to incubate at room temperature or at 37°C for a minimum of 2 hours. On day 0, H1 ESCs were detached applying Accutase (Sigma-Aldrich), previously heated to 37°C, for 4 minutes at 37 °C, 5% CO_2_. Following detachment, cells were diluted in preheated DMEM/F-12, dissociated into single cells by repeated pipetting and transferred into a 50 mL canonical tube. ESCs were pelleted by centrifugation at 300 g for 8 minutes, at room temperature. Subsequently, the supernatant was removed by aspiration and cells resuspended in STEMdiff Neural Induction Medium supplemented with STEMdiff SMADi Neural Induction Supplement and 10 μM Rho-associated protein kinase (ROCK) inhibitor Y-27632 (Stem Cell Technologies). The obtained single cell suspension was counted, and, following removal of the coating solution, cells were distributed into the previously prepared plates at a density of 2-2.5 x 10^5^ cells/cm^2^. Daily medium changes were performed.

Cells were passaged weekly and maintained in STEMdiff Neural Induction Medium supplemented with STEMdiff SMADi Neural Induction until passage 3. Following passage 3, cells were maintained in STEMdiff Neural Progenitor Basal Medium (Stem Cell Technologies). For passaging, cells were detached by incubation in 1 mL of pre-heated Accutase for 8 minutes. Aggregates of cells were dissociated using a 1 mL pipette and resuspended in 3 mL of DMEM/F-12, pre-wared to 37°C. Wells were subsequently transferred into a 50 mL canonical tube, and further diluted with DMEM/F-12. NPCs were pelleted through centrifugation at 300 g for 5 minutes, at room temperature. The supernatant was subsequently removed, and cells resuspended in the according medium. To eliminate cell clumps, prone to promote spontaneous differentiation, cells were filtered through a 40 μm cell strainer. Cells were subsequently counted and plated at the desired density in the previously coated plates. For cryopreservation, cells were instead counted prior to pelleting, and, following centrifugation, resuspended in STEMdiff Neural Progenitor Freezing Medium (Stem Cell Technologies) to obtain a final density of 2 x 10^6^ cells/mL. The NPCs were distributed within cryotubes, transferred into a Mr. Frosty Freezing Container (Thermo Scientific) and gently frozen at -80°C. For long-term storage, cells were transferred to -150°C.

### Generation and maintenance of human forebrain neurons

Human forebrain neurons were generated starting from the previously obtained NPCs (see “Generation and maintenance of human neural progenitor cells”) using the STEMdiff Forebrain Neuron Differentiation Kit (Stem Cell Technologies) and the STEMdiff Forebrain Neuron Maturation Kit (Stem Cell Technologies) as per manufacturer’s recommendations. Briefly, 6-well plates or 24-well μ-plates were coated as previously described (see “Generation and maintenance of human neural progenitor cells”), using a lower concentration of laminin of 5 μg/mL. All plates were pre-warmed to 37°C before use. On day 0, NPCs were passaged as aforementioned. Cells were subsequently resuspended in an appropriate volume of STEMdiff Neural Induction Medium or STEMdiff Neural Progenitor Basal Medium, depending on the NPC passage. Cells were seeded at a density of 125’000 – 140’000 cells/cm^2^ and left to incubate at 37°C, 5% CO_2_ overnight. On the following day, cell culture medium was replaced with STEMdiff Forebrain Neuron Differentiation Medium, prepared as per manufacturer’s instructions. Neuronal precursor cells were maintained with daily medium changes until a confluency on 80-90% was confirmed. Once confluency was observed, STEMdiff Forebrain Neuron Differentiation Medium was aspirated, 1 mL of Accutase was added to each well and cells left to detach at 37°C, 5% CO_2_ for 8 minutes. Cells were resuspended in 3 mL of pre-heated DMEM/F-12 and transferred into a 50 mL conical tube. Neuronal precursor cells were further washed by adding DMEM/F-12 to reach 50 mL and subsequently pelleted by centrifugation at 400 g for 5 minutes. The supernatant was discarded, cells were resuspended in an appropriate volume of STEMdiff Forebrain Neuron Maturation Medium and counted. Neuronal precursors were seeded onto pre-heated coated cultureware at a density of 4.8 x 10^4^ – 6.6 x 10^4^ cells/cm^2^. Full medium changes were performed in 2–3-day intervals and cells were cultured for a minimum of 8 days before infection.

### Vero E6 culture maintenance

Vero E6 cells (kindly provided by Dr. Doreen Muth, Dr. Marcel Müller, and Dr. Christian Drosten, Charité Berlin, Germany) were cultured at 37°C, 5% CO_2_ in DMEM supplemented with GlutaMAX (Gibco), 10% fetal bovine serum (FBS, Gibco), 1 mM Sodium Pyruvate (Gibco) and 10 mM HEPES (Gibco). Cells were passaged twice per week and used up to a maximum number of 20 passages. For MPXV infection, cells were seeded in 25 cm^2^ tissue culture flasks (TPP), at a density of 1.4 x 10^6^ cells per flask with addition of 1% Penicillin-Streptomycin (Sigma-Aldrich, Gibco), and cultured overnight at 37°C, 5% CO_2_. Medium was allowed to reach room temperature prior to application on cells.

### Generation of human neural organoids

hNOs were generated using an adapted version of the previously published protocol by Lancaster and colleagues (2014)^46^. Briefly, on day 0, stem cells were detached applying Accutase for 4 minutes at 37 °C, 5% CO_2_. Cells were dissociated by repeated pipetting till a single-cell suspension was obtained. Cells were counted, centrifuged at 200 g for 5 minutes at 4°C, and resuspended in the required volume of Formation Medium prepared using the STEMdiff Cerebral Organoid Kit (Stem Cell Technologies) supplemented with 50 μM Y-27632. The cell suspension was distributed into ultra-low attachment 96-well plates (Corning), each well containing 9’000 to 10’000 cells suspended in 100 μL of Formation Medium. The cultures were left undisturbed for at least 24 hours. On day 2 and 4, 100 μL/well of Formation Medium without ROCK inhibitor were added. On day 5, the formation of embryoid bodies (EBs) was confirmed and EBs were transferred with wide-bore 200 μL tips to ultra-low attachment 24-well plates (Corning), previously filled with 500 μL/well of Induction Medium, prepared using the STEMdiff Cerebral Organoid Kit. On day 7, EBs were embedded into 30 μL droplets of Matrigel (Corning) each and transferred into ultra-low attachment 6-well plates (Corning) containing 3 mL of home-made Expansion Medium consisting of a 1:1 mixture of DMEM/F-12 and Neurobasal Medium (Gibco) supplemented with 1:200 N2 (Gibco), 1:100 B27 minus vitamin A (Gibco), 2.5 μg/mL insulin (Sigma-Aldrich), 1:100 GlutaMAX (Gibco), 1:200 MEM-NEAA (Seraglob), 3.5 μL/L 2-mercaptoethanol (Sigma-Aldrich), and 1:100 Penicillin-Streptomycin. On day 10, Expansion Medium was gently removed and replaced with 3.5 mL homemade differentiation medium consisting of a 1:1 mixture of DMEM/F-12 and Neurobasal medium supplemented with 1:200 N2, 1:100 B27, 2.5 μg/mL insulin, 1:100 GlutaMAX, 1:200 MEM-NEAA, 3.5 μL/L 2-mercaptoethanol, 1:100 Penicillin-Streptomycin, and 0.5 μg/mL Amphotericin B (Gibco). From day 10 on, hNOs were cultured on an orbital shaker at 65 rpm, in a 37 °C, 5% CO_2_ incubator. Medium was changed every 2 to 3 days. hNOs between 70-110 days of culture that passed the quality control criteria^46^ were used for the experiments. Criteria included ESC and iPSC stemness of at least 80% of double positive cells for both stemness markers SSEA-5 (Stem Cell Technologies, 60063AZ, 1:50) and TRA-1-60 (Stem Cell Technologies, 60064PE, 1:50), which was measured at day 0 by flow cytometry (FCM) assay. For this, a sample of cells equivalent to 10^6^ cells was pelleted by centrifugation at 280 g for 8 minutes at 4°C. Following centrifugation, the supernatant was removed by aspiration and cells were resuspended in 100 μL of BD CellWASH (BD) supplemented with the previously mentioned antibodies. Cells were thoroughly vortexed, incubated at 4°C in the dark for 15 minutes and subsequently washed through addition of 1 mL of BD CellWash, followed by centrifugation at 280 g for 8 minutes at 4°C. The supernatant was removed once more, and cells resuspended in 100 μL of BD CellWASH. FCM acquisitions were performed on a FACS Canto II (BD Bioscience) using the DIVA software and further analyzed with FlowJo (TreeStar). Further quality control criteria included typical sizes corresponding to each developing stage, brightening of outer layer at the EBs stage before embedding, and formation of neural tube-like structures in Matrigel. All used media were warmed to room temperature before usage.

### Monkeypox virus propagation

A primary isolate of MPXV belonging to clade IIb lineage isolated in 2022 was used for this study (kindly provided by Prof. Dr. Isabela Eckerle, Geneva Center for Emerging Viral Diseases and Division of Infectious Diseases, University Hospital of Geneva). The MPXV isolate was passaged three times on Vero E6 cells to produce virus stocks. Briefly, 24 hours prior to infection, Vero E6 cells were seeded in 75 cm^2^ tissue culture flasks (TPP), at a density of 8 x 10^6^ cells per flask, and cultured at 37°C, 5% CO_2_ in the previously described medium. On the day of infection, the culture medium was replaced with 5 mL of DMEM supplemented with GlutaMAX and 2% FBS, and the required amount of virus stock was added. Cells were placed on an orbital shaker moving at 60 rpm for 1 hour at 37°C, 5% CO_2_. Following incubation, 15 mL DMEM supplemented with GlutaMAX and 2% fetal bovine serum were added. Cells were incubated at 37°C, 5% CO_2_ and inspected daily for signs of cytopathic effect (CPE). At day 3 p.i. (i.e., when a CPE of around 80% of cells was observed) flasks were frozen at -70°C, to release cell-associated viral particles. Viral suspensions were subsequently thawed, collected in 50 mL conical tubes, and centrifuged at 1000 g for 10 minutes at 4°C to remove cell debris. Supernatants were aliquoted and stored at -70°C till further use. Mock control stocks were generated in parallel, following the same procedures, with omission of virus inoculum. All procedures involving MPXV handling were carried out in biosafety level 3 (BSL3) conditions at the Institute of Virology and Immunology (University of Bern, Switzerland) by trained personnel provided with appropriate personal protective equipment. All media and reagents were warmed to reach room temperature prior to application on cells.

### Construction of MPXV expressing mNeonGreen

For the construction of an mNG-expressing MPXV, the transfer vector pUC57-MPXV-GPT-mNG carrying a DNA cassette with vaccinia virus promoter-driven *E. coli* Xanthine-guanine phosphoribosyltransferase (GPT)^125^ and mNG^126^ genes flanked by partial MPXV I8R and G1L gene sequences (Supplementary Figure 3h) was generated by oligonucleotide synthesis (GenScript). The cassette encodes *E. coli* GPT under the control of the synthetic vaccinia virus promoter (P-VV syn (E/L) obtained from the GenBank entry ON549927.1 (Mutant Cowpox virus strain recombinant Brighton Red clone deltaCPXV014). The GPT cassette was followed by the mNG reporter gene under the early-late vaccinia virus VV7.5 promoter (P-VV P7.5) sequence obtained from the GenBank sequence GU062789.1 (Poxvirus recombinant transfer vector pSPV-EGFP). For recombination-based integration of the GPT-mNG cassette into the intergenic region between I8R (OPG084) and G1L (OPG085) of MPXV, the GPT-mNG cassette was edged by the corresponding flanking regions, each 700 base pairs in length, similar to a report by Freyn A.W. and colleagues^127^. The plasmid construct was amplified using One Shot Stbl3 Chemically Competent *E. coli* cells (Thermo Fisher Scientific). To determine the identify and the purity of recombinant MPXV, DNA fragments corresponding to the OPG085-OPG084 locus were amplified from viral genomic DNA purified with the NucleoSpin Virus Kit (Macherey-Nagel) using the Phusion Hot Start II DNA Polymerase (Thermo Fisher Scientific) and specific pairs of oligonucleotides OPG085-ext-F1 (GCGACGATTATATGATGAAC) and OPG084-ext-R1 (ACATATTCCTGGAGATACAC) or OPG085-ext-F2 (CCTTGGCTATCGCATGATTT) and OPG084-ext-R2

(CATCCGGTATAGTCTTTGTG) (Microsynth). The recombinant virus was generated by infection of Vero E6 cells with the parental MPXV at an MOI of 1 TCID_50_/cell followed by transfection of 1 μg of transfer vector with Lipofectamine 2000 reagent (Thermo Fisher Scientific) in OptiMEM (Gibco) following manufacturer’s recommendations. The recombinant virus was purified by four rounds of isolation of fluorescent plaques under GPT selection (Merck).

### Monkeypox virus titration

Viral stocks, cell culture and organoid supernatants as well as homogenized cells and organoids were titrated on Vero E6 cells in a 96-well plate format. 24 hours prior to titration, the required number of 96 well plates (TPP) were seeded with a suspension of Vero E6 cells at a density of 2 x 10^4^ cells per well. Serial dilutions were performed in DMEM supplemented with GlutaMAX, 2-10% FBS,1 mM Sodium Pyruvate, 10 mM HEPES (Gibco), and 1% Penicillin-Streptomycin, starting at a dilution of 1:10. Cell supernatant was aspirated, and the dilutions transferred on the previously seeded cells. Cells were incubated for 72 hours at 37°C, 5% CO_2_. For readout, the inoculum was aspirated, cells washed once with 200 μL DPBS (Sigma-Aldrich), fixed for 15 minutes by immersion in 4% formalin (Formafix), and subsequently stained for 10 minutes with crystal violet. Plates were washed with tap water and left to dry prior to readout. Viral titers were calculated as TCID_50_ per mL using the Reed and Muench method. For titration of released viral loads, culture supernatant was collected at selected time points and stored at -70°C until further analysis. Analogously, for quantification of cell-associated infectious virus, one organoid per time point was randomly sampled, washed with sterile DPBS and transferred into a 1.5 mL screw cap tube containing 1 mL of sterile DPBS. The sample was frozen at -70°C. For analysis, the thawed hNO was dissociated thoroughly by pipetting up and down, until no cell clumps were visible any longer. The sample was centrifuged at 450 g for 5 minutes at 4°C and the supernatant used for titration. For titration of cell-associated live MPXV in 2D neural cultures, cells were washed three times with sterile DPBS, topped with 1 mL of sterile DPBS and frozen at -70°C until analysis. Vero E6 cells were washed twice with 5 mL of sterile DPBS, topped with 2 mL of sterile DPBS and stored at - 70°C. For analysis, cells were thawed and subsequently scratched off the cultureware using a cell scratcher or inverted 1 mL pipette tip. Samples were subsequently centrifuged at 450 g for 5 minutes at 4°C, and the supernatant used for quantification. For normalization of hNO titers, obtained released viral titers were multiplied by the volume of medium per well and divided by the number of hNOs per well, to obtain released infectious virus per hNO. Analogously, cell-associated viral titers were multiplied by the total sample volume and divided by the number of organoids per sample, to obtain cell-associated infectious virus per hNO. For normalization of 2D cell culture titers, obtained released viral loads were multiplied by the total volume of medium per well or flask. Analogously, cell-associated viral titers were multiplied by the total sample volume, to obtain total infectious virus content. Medium and reagents were warmed to room temperature prior to application on cells.

### Neural progenitor cell infection and reporter construct fluorescence imaging

NPCs were infected one day following passaging in 24 or 6 well plates with an MOI of 0.01 TCID_50_/cell. Considering the growth rate anticipated as per manufacturer’s instructions, for MOI calculation, no doubling was considered within the first 24 hours post-seeding. Virus suspension was prepared by adding the calculated volume of virus stock to the required amount of STEMdiff Neural Progenitor Basal Medium in a 50 mL conical tube to obtain a homogeneous suspension of 500 μL or 2 mL per well, depending on plate format. Mock mixture was prepared analogously. The NPC medium was removed from all wells and substituted with inoculum. Cells were subsequently incubated for 2 hours at 37°C, 5%CO_2_. Following incubation, the inoculum was aspirated, and cells were washed three times with 500 μL or 3 mL of sterile DPBS. After washing, 500 μL or 2 mL of STEMdiff Neural Progenitor Basal Medium were distributed into each well, and NPCs were placed back in the incubator at 37°C, 5% CO_2_. Half-volume medium changes were performed daily. Supernatant samples were collected at predefined time points and stored at -70°C till analysis. For cell-associated virus titration, the medium was removed and wells washed three times with 3 mL of sterile DPBS. Following the last washing steps, 1 mL of sterile DPBS was added into each well, and plates were subsequently frozen at -70°C till further analysis. Light microscopy images were taken using an EVOS XL or EVOS M7000 Imaging System (Thermo Fisher Scientific). For fluorescence imaging and time-lapse live-cell microscopy, NPCs challenged with an MPXV clade IIb lineage mNG reporter strain were imaged at regular intervals using an EVOS M7000 Imaging System with EVOS Onstage Incubator. Brightness and contrast of all micrographs was adjusted using ImageJ. All media and reagents were allowed to reach room temperature prior to application on cells.

### Forebrain neuron infection and reporter construct fluorescence imaging

Forebrain neuron cultures were infected after a minimum maturation time of 8 days in 24 or 6 well plates at a rate of 0.01 TCID_50_/cell. For accurate MOI calculations, 24 hours pre-infection cells of one well were detached, dissociated and counted to obtain total neuron numbers per well. MPXV inoculum was obtained by adding the calculated volume of virus stock to a predefined amount of STEMdiff Forebrain Neuron Maturation Medium, corresponding to 500 μL or 2 mL of medium per well. The suspension was thoroughly mixed and distributed upon the cells, after cell culture medium aspiration. Infection was allowed to occur for 2 hours at 37°C, 5% CO_2_. The virus suspension was subsequently removed by aspiration, and cells were gently washed three times with 500 μL or 3 mL of sterile DPBS. Following the last washing step, 500 μL or 2 mL of STEMdiff Forebrain Neuron Maturation Medium were added, and neurons were placed back into the incubator at 37°C, 5% CO_2_. Supernatant samples were collected at predetermined time points and stored at -70°C until analysis. For cell-associated virus quantification, cell culture medium was discarded, and neurons were washed three times with 3 mL of sterile DPBS. 1 mL of sterile DPBS was subsequently added on top of the neurons and plates were stored at -70°C until analysis. Light microscopy images were taken at regular intervals using an EVOS XL or EVOS M7000 Imaging System. Neurons challenged with MPXV clade IIb lineage mNG reporter construct were either imaged at regular intervals or continuously through time-lapse live-cell imaging, using an EVOS M7000 Imaging System with EVOS Onstage Incubator. Brightness and contrast of all micrographs was adjusted using ImageJ. All media and reagents were allowed to reach room temperature prior to application on cells.

### Vero E6 infection

Vero E6 cells were infected one day following passaging in 25 cm^2^ tissue culture flasks with an MOI of 0.01 TCID_50_/cell. For MOI calculation, one cell doubling per 24 hours was considered. Virus mixture was prepared by adding the calculated volume of virus stock to DMEM supplemented with GlutaMAX, 2% FBS,1 mM Sodium Pyruvate, 10 mM HEPES and 1% Penicillin-Streptomycin into a 50 mL conical tube to obtain a homogeneous suspension of 5 mL per flask. Mock suspension was prepared analogously. The cell culture medium was removed from all flasks and substituted with the inoculum. Infection was allowed to proceed for 2 hours at 37°C, 5% CO_2_. The inoculum was subsequently removed and cells washed three times with 5 mL of sterile DPBS. 6 mL of the previously described medium were subsequently added to each flask, and cells placed back into the incubator at 37°C, 5% CO_2_. Supernatant samples were collected in a daily basis for a total of 4 days p.i and stored at -70°C until analysis. For cell-associated live virus quantification, cell culture medium was removed from the flasks completely and cells were washed twice with 5 mL of sterile DPBS. 2 mL of DPBS were added, and cells were frozen at -70°C until analysis. Light microscopy images were taken at predefined timepoints using an EVOS XL Imaging System. Brightness and contrast of all micrographs was adjusted using ImageJ. All media and reagents were allowed to reach room temperature prior to application on cells.

### Human neural organoids infection

hNOs were infected at 70 to 110 days of development in a 6-well plate format with an MOI of 0.1 TCID_50_/cell. MOI calculations were based on previously determined average cell numbers per hNO, derived from routine cell counts of dissociated organoids using FCM at various developmental stages. Briefly, virus suspension was prepared by adding the required amount of virus stock to organoid Maturation Medium (see “Generation of human neural organoids”) in a 50 mL conical tube to obtain a homogeneous suspension of 3 mL per well. Mock treatment was prepared analogously. Organoid culture medium was removed from all wells and replaced with the inoculum. hNOs were incubated for 2 hours at 37°C, 5% CO_2_, constantly shaking at 65 rpm. hNOs were subsequently washed three times with 3 mL of sterile DPBS. 4 mL of fresh organoid Maturation Medium were added to each well, and hNOs were placed back in the incubator at 37°C, 5% CO_2_, shaking at 65 rpm. Medium changes were performed in two-day intervals. Tissue and culture supernatant samples were collected at selected time points and stored at -70°C until further analysis. Light microscopy images were taken at regular intervals using an EVOS XL Imaging System. Images were processed using ImageJ for brightness and contrast optimization and Imaris 9.9.1. (Bitplane) for surface area measurements. All media and reagents were allowed to reach room temperature prior to application of hNOs.

### Human neural organoid tecovirimat treatment

For tecovirimat treatment, hNOs were infected as described above (“Human neural organoids infection”). Following washing, organoid Maturation Medium was supplemented with tecovirimat (Selleckchem) at predefined concentrations. Medium changes were performed in two-day intervals with tecovirimat supplemented medium. Tissue and culture supernatant samples were collected at selected time points and stored at -70°C until further analysis. Medium and reagents were allowed to reach room temperature prior to application of hNOs.

### 2D human neural cell immunofluorescence

For confocal analysis of 2D neural cultures, cells seeded onto 24-well μ-plates as previously described (see “Generation and maintenance of human neural progenitor cells” and “Generation and maintenance of human forebrain neurons”) were used. Cells were fixed in 4% formalin and subsequently washed and permeabilized for 10 minutes in a mixture of 0.1% Tween-20 (Sigma-Aldrich) in PBS. To prevent unspecific staining, cells were subsequently blocked with a 5% goat serum (Sigma-Aldrich) in 0.1% Tween-20 solution for 1 hour at room temperature. Primary antibody solutions were prepared in 0.1% Tween-20 solution supplemented with either 5% bovine serum albumin (Sigma-Aldrich) or 5% goat serum at the following dilutions: TUJ1 (BioLegend, 801202, 1:400), SOX2 (Invitrogen, 14-9811-82, 1:200). Following incubation, blocking buffer was gently removed and substituted with 500 μL of primary antibody solution per well. Plates were sealed with PARAFILM M (Sigma-Aldrich) and incubated over night at 4°C. On the following day, cells were washed three times for 5 minutes with 500 μL of 0.1% Tween-20 in PBS solution to ensure complete antibody removal. Secondary antibody mixtures were prepared in 0.1% Tween-20 solution at the following dilutions: anti-mouse IgG2a AF594 (Invitrogen, A21135, 1:200), anti-rat AF647 (Invitrogen, A21247, 1:500). For nuclear staining, DAPI (Sigma-Aldrich, 62248, 1:600) was added to the secondary antibody solution. 500 μL of secondary antibody mixture were distributed into each well and staining was allowed to proceed for 1.5 hours at room temperature, in the dark. After staining completion, wells were washed three more times for 5 minutes with 500 μL of 0.1% Tween-20 in PBS solution and finally topped with 500 μL of DPBS. Imaging was performed using a Carl Zeiss LSM710 confocal microscope at the microscopy imaging center (MIC) of the University of Bern, Switzerland. Brightness and contrast of micrographs was adjusted using ImageJ.

### Human neural organoids cryopreservation and immunofluorescence

hNOs were processed using an adapted version of the protocol “Cryogenic Tissue Processing and Section Immunofluorescence of Cerebral Organoids” provided by Stem Cell Technologies. For immunofluorescence, hNOs were washed twice with PBS and fixed in 4% formalin. Next, hNOs were washed three times with PBS. For cryogenic preservation, hNOs were subsequently incubated at 4°C in a 30% sucrose (Sigma-Aldrich) in PBS solution until organoids were observed to sink to the bottom of the tube. Equilibrated hNOs were embedded and incubated in a 7.5% gelatin, 10% sucrose in PBS solution for 1 hour at 37°C, to avoid polymerization. In the meantime, a mixture of dry ice and 100% ethanol was prepared. hNOs and gelatin were subsequently transferred to a Cryomold (Biosystems Switzerland AG) and, after initial polymerization at room temperature, carefully placed on the surface of the (no longer boiling) mixture of dry ice and ethanol. Once completely frozen, embedded hNOs were transferred to -80°C for long term storage. For immunofluorescence, cryopreserved hNOs were processed into 18 µm thin slices using a Leica CM1950 cryostat and placed on SuperFrost Plus Adhesion slides (Epredia). Slides were stored at -20°C till staining. For staining, sections were allowed to reach room temperature prior to processing. To dissolve gelatin, slides were subsequently incubated for 10 minutes in a 37°C pre-warmed mixture of 0.1% Tween-20 in PBS. hNO sections were encircled using a ReadyProbes Hyrophobic Barrier Pap Pen (Thermo Fisher Scientific) and blocked with a 5% donkey serum (Abcam) or 5% goat serum in 0.1% Tween-20 in PBS solution for 1 hour at room temperature. Primary antibodies were resuspended in 0.1% Tween-20 in PBS supplemented with 5% bovine serum albumin at the following dilutions: TUJ1 (BioLegend, 801202, 1:400), SOX2 (Invitrogen, 14-9811-82, 1:200), GFAP (Invitrogen, PA1-10004, 1:200), vaccina virus i.e. MPXV A27L protein (OriGene Technologies, Inc BP1076, 1:1000), cleaved caspase-3 (arigo Biolaboratories Crop., ARG66888, 1:200). Blocking buffer was removed, 100 μL of primary antibody solution were added to each section and slides were allowed to incubate at 4°C overnight. On the following day, primary antibody solution was removed by gently tapping slides on paper towels and sections were washed three times for 5 minutes by immersion in 0.1% Tween-20 in PBS. Secondary antibodies were resuspended in 0.1% Tween-20 in PBS at the following dilutions: anti-mouse AF488 (Invitrogen, A21131, 1:200), anti-mouse AF647 (Invitrogen, A2124, 1:200), anti-rat AF647 (Invitrogen, A21247, 1:500), anti-chicken AF546 (Invitrogen, A11040, 1:200), anti-rabbit AF488 (Jackson ImmunoResearch Europe Ltd., 711-545-152, 1:200), anti-rabbit AF546 (Invitrogen, A11035, 1:200), anti-rabbit AF647 (Invitrogen, A21245, 1:200). For nuclear and F-actin staining, DAPI (Sigma-Aldrich, 62248, 1:1000) and Rhodamine Phalloidin (Invitrogen, R415, 1:200) were added to the mixture. Sections were incubated with secondary antibodies for 2 hours at room temperature. Slides were subsequently washed three more times for 5 minutes by immersion in 0.1% Tween-20 in PBS and mounted in EMS Shield Mount with DABCO (Electron Microscopy Sciences). Imaging was performed using a Carl Zeiss LSM710 confocal microscope at the microscopy imaging center (MIC) of the University of Bern, Switzerland. Brightness and contrast of micrographs was adjusted equally between groups using ImageJ. For complete visualization of beaded filaments, multiple z-stack planes were optimized and merged.

### Transmission electron microscopy

For transmission electron microscopy, hNOs were fixed using a 2.5% glutaraldehyde and 2% formalin solution in organoid Maturation Medium for a duration of 1 day at 4°C. Subsequently, the fixation solution was replaced with 4% formalin, and samples stored at 4°C until further processing. hNOs were cut into 1 mm3 cubes and immerged in a 1% osmium tetroxide (Electron Microscopy Sciences) solution in ddH_2_O for 1 hour at room temperature. After washing three times with ddH_2_O, samples were progressively dehydrated in a graded ethanol series, starting from 50% and progressing to 70%, 80%, 90%, and finally 100% ethanol. The samples were then embedded in epoxy resin (EM0300, Sigma-Aldrich) and left to polymerize for 8 hours at 70℃. Ultrathin sections of resin-embedded tissue were prepared using a Leica EM UC7 ultramicrotome. Sections of 70 nm were collected on EM grids (G100H-Cu, Electron Microscopy Sciences) and processed using a 2% uranyl acetate and 0.5% lead citrate solution for contrast enhancement. The obtained sections were imaged using a CM-100 transmission electron microscope (Philips) at 80 kV. Brightness and contrast of all micrographs was adjusted using ImageJ.

### Human neural organoid RNA isolation

For RNA isolation from hNOs, organoids were washed in sterile DPBS before being transferred into MagNA Lyser Green Beads tubes (Roche) containing 5 ceramic beads per tube and 700 μL of TRIzol reagent (Thermo Fisher Scientific) cooled to 4°C. hNOs were shredded using a Tissue Homogenizer Bullet Blender BBX24B-CE (Next Advance) at speed 8 for 1 minute. Alternatively, 50 μL of hNO homogenate in DPBS were added to 700 μL of TRIzol reagent. Tubes were subsequently stored at -70°C until further analysis. For RNA isolation, TRIzol homogenates were thawed on ice, 0.2 mL of chloroform (Sigma Aldrich) were added and samples incubated on ice for 5 minutes. Samples were precipitated at 12’000 g for 15 minutes at 4°C. The obtained RNA-containing aqueous phase was transferred to a new tube and supplemented with an equal volume of absolute ethanol (Sigma Aldrich). Subsequent purification was performed on Zymo-Spin IC Columns (Zymo Research) as per manufacturer’s recommendations. Purified RNA was eluted in 40 μL of RNAse-free water. RNA quality was evaluated using a Fragment Analyzer (Agilent) and ProSize Data Analysis Software (Agilent).

### Monkeypox virus DNA isolation

MPXV virus DNA was isolated from hNOs using the QIAamp DNA Blood Mini Kit (QIAGEN). For sample lysis and virus inactivation, 200 μL of supernatant (for extracellular MPXV DNA extraction) or 200 μL of hNO homogenate in DPBS (for cell-associated virus quantification) were added to 200 μL of Buffer AL within 1.5 mL tubes and thoroughly mixed by repeated pipetting. For cell-associated virus DNA extraction, 20 μL of reconstituted QIAGEN Protease were added to the mixture and the solution was pulse-vortexed for 15 seconds. The sample was then spun down and incubated at 56°C for 10 minutes. After a second spin, the obtained mixture was transferred to a new 1.5 mL tube and 200 μL of absolute ethanol were added. The sample was once more thoroughly mixed by pulse-vortexing for 15 seconds, spun down and stored at -70°C until further analysis. DNA was subsequently extracted by applying the obtained solution to QIAamp Mini spin columns and processed as per manufacturer’s instructions. Isolated DNA was eluted in 200 μL of Buffer AE.

### Real-time quantitative PCR

For inflammatory response analysis, the Omniscript RT Kit (QIAGEN) with Random Primers (Invitrogen) was used for complementary DNA (cDNA) synthesis from previously isolated hNO RNA (see “Human neural organoid RNA isolation”). MPXV DNA, previously isolated from cell culture supernatants and lysed hNOs, was used for viral load quantification through qPCR (see “Monkeypox virus DNA isolation”). A PCR Master Mix containing RNase-free water (QIAGEN), TaqMan Fast Universal PCR Master Mix (2X), no AmpErase UNG (Applied Biosystems), target-specific forward and reverse primers and probe was prepared. Two μL of the previously obtained cDNA or previously isolated MPXV DNA were added to 18 μL of master mix for each reaction and processed using an Applied Biosystems 7500 Real-Time PCR System (Applied Biosystems). Released viral loads were normalized as gene copies per mL. Cell-associated viral loads, cytokine and chemokine relative expression was calculated using the previously described ΔΔCT approach^128^, and normalized to the expression of the housekeeping 18S rRNA. Previously published primer and probe sequences were used to detect MPXV DNA^129^, IFN-β^130^, IL-6^131^, IL-8/CXCL8 and IP-10/CXCL10^132^.

### RNA sequencing analysis

Bulk RNA barcoding and sequencing (BRB-Seq) libraries^133^ were generated as per manufacturer’s recommendation (Alithea Genomics). Libraries were then processed as described in the publication above. Briefly, the FASTQ files were generated using the Illumina software bcl2fastq (v2.20.0.422) and were mapped to the human reference genome (GRCh38, annotation: Ensembl GRCh38.111) using STAR^134^ (v. 2.7.10b). 411,101,117 reads (82.16%) were uniquely mapped to the genome. Analysis of gene expression was made with DESeq2^135^ (v1.42.0). The DESeq2 design took into account day and infection and corrected for biases produced by row/column positioning in the BRB-seq oligo(dT) primer plate. In the differential expression analyses, genes were differentially expressed if they had an adjusted p-value equal or lower than 0.05 between libraries. Gene ontology analyses were performed with ClusterProfiler^136^ (v4.10.0). GO terms were considered significant if they had an adjusted p-value equal or lower than 0.05.

### Statistical analysis

GraphPad Prism 9 software was used for statistical analysis and graph generation. Two-sided t-tests or non-parametrical Mann-Whitney U tests were used to assess significance of detected values. p < 0.05 was considered statistically significant.

## ACKNOWLEDGEMENTS

We would like to extend our gratitude to the Geneva Center for Emerging Viral Diseases, the Division of Infectious Diseases and Prof. Dr. Isabela Eckerle for providing the clade IIb lineage MPXV isolate used in this study. This work was supported by a grant from the Multidisciplinary Center for Infectious Diseases of the University of Bern (to M.P.A.) and by intramural funding of the Institute of Virology and Immunology. Illustrations were created using BioRender: Schultz-pernice, I. (2025) https://BioRender.com/o02y129.

## DATA AVAILABILITY STATEMENT

The data generated and analyzed during the current study are available from the corresponding author on reasonable request.

## AUTHOR CONTRIBUTIONS

I.S.P., A.F., D.B., and M.P.A. conceptualized and designed the project. I.S.P., A.F., F.B., M.L., Y.C.C., B.O.I.E., T.D., and B.Z. conducted the investigations. I.S.P., A.F., F.B., M. L., Y.C.C., B.O.I.E., T.D., B.Z., A.G., M.G., S.S., C.W., R.Z. D.J., F.B., and O.B.E., A.S., and N.R developed the methodology and provided reagents. M.P.A., I.S.P., A.F., F.B., M.L., Y.C.C., B.O.I.E., T.D., and M.G. curated and analyzed the data. I.S.P., A.F., and M.P.A. wrote the original manuscript. All authors reviewed, edited, and approved the manuscript.

---

**Supplementary Figure 1:**
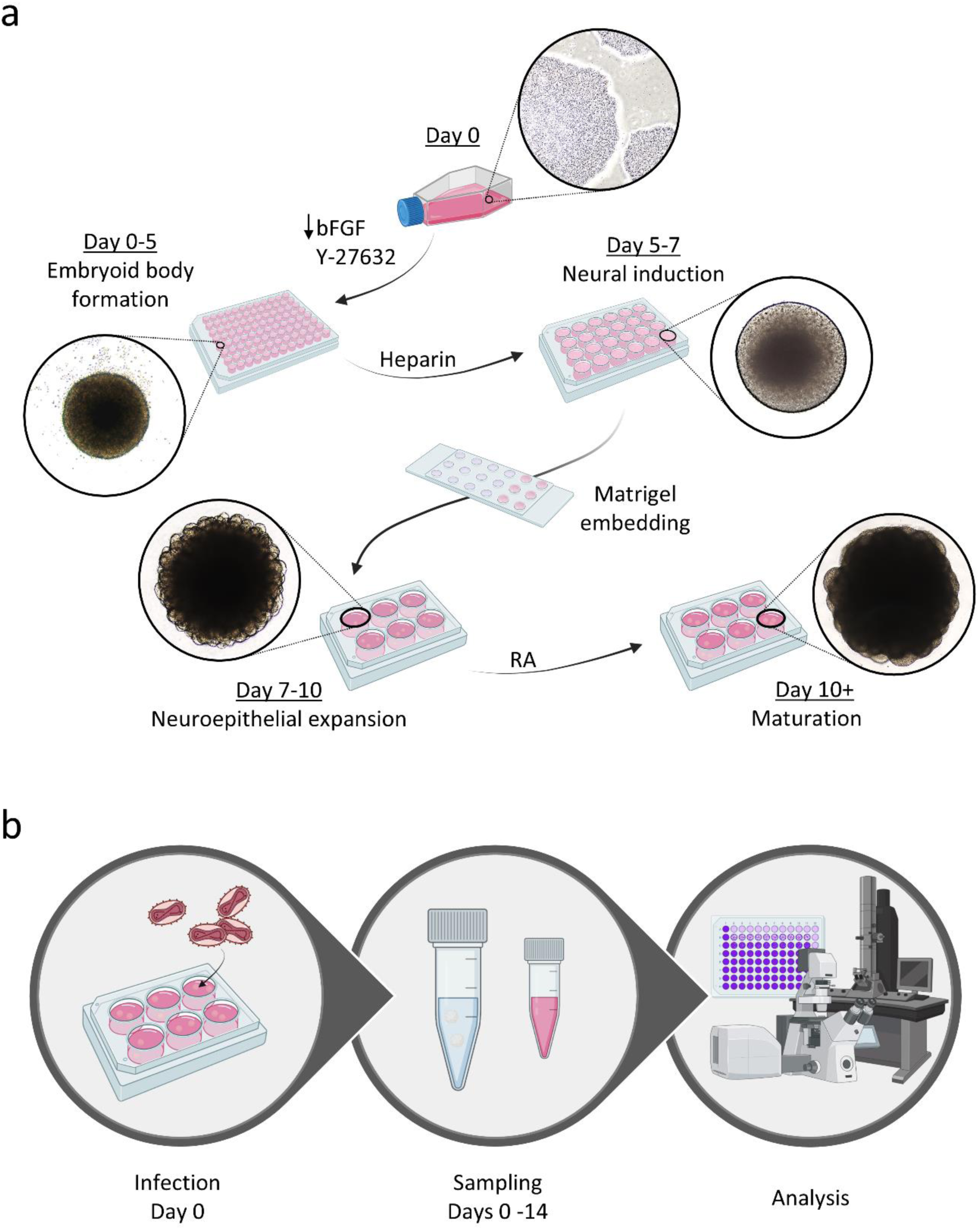
Experimental design for hNOs generation and infection. (a) At day 0 pluripotent stem cells were dissociated, resuspended in Formation Medium and distributed at a density of 9’000-10’000 cells per well in ultra-low attachment 96-well plates for EB formation. After 5 days, EBs were transferred to Induction Medium for neural induction. At day 7, EBs were embedded into Matrigel and transferred to ultra-low attachment 6-well plates in Expansion Medium. At day 10, hNOs were transferred to Maturation Medium and moved to an orbital shaker for further development. (b) After 70 to 110 days of culture, hNOs were infected after 70 to 110 days of culture with a clade IIb lineage MPXV isolate at an MOI of 0.1 TCID_50_/cell. Supernatant and tissue samples were taken at regular intervals for a period of up to 14 days, prior to analysis.

**Supplementary Figure 2:**
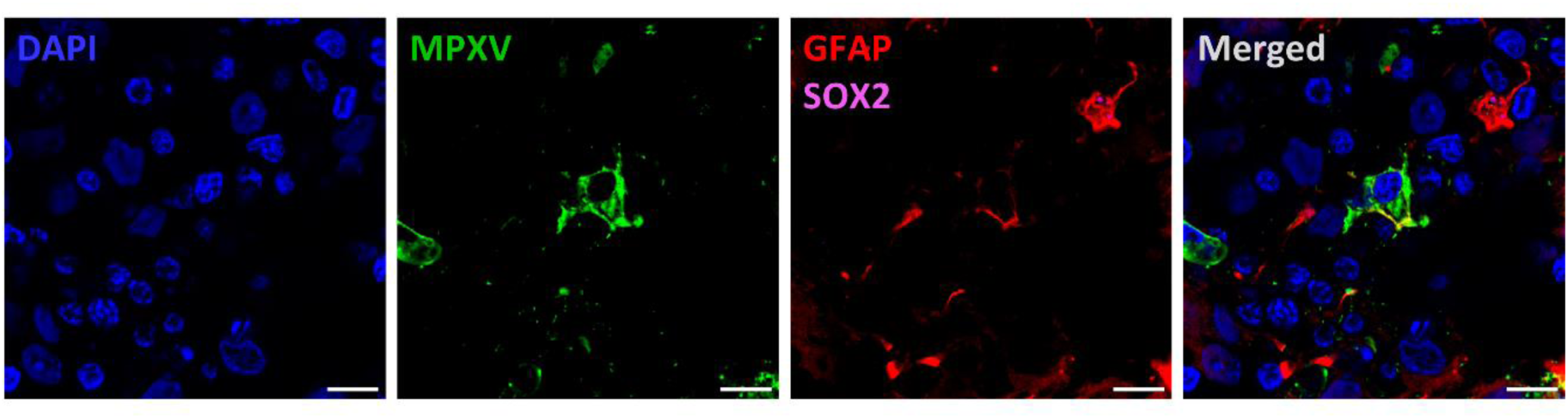
Astrocytes are susceptible to MPXV infection in hNOs. Representative micrographs displaying MPXV tropism for astrocytes in 70-75 days old hNOs 10 days p.i. with a clade IIb lineage isolate at an MOI of 0.1 TCID_50_/cell. DAPI, blue; MPXV, green; SOX2, pink; GFAP, red. Scale bar, 10 µm.

**Supplementary Figure 3:**
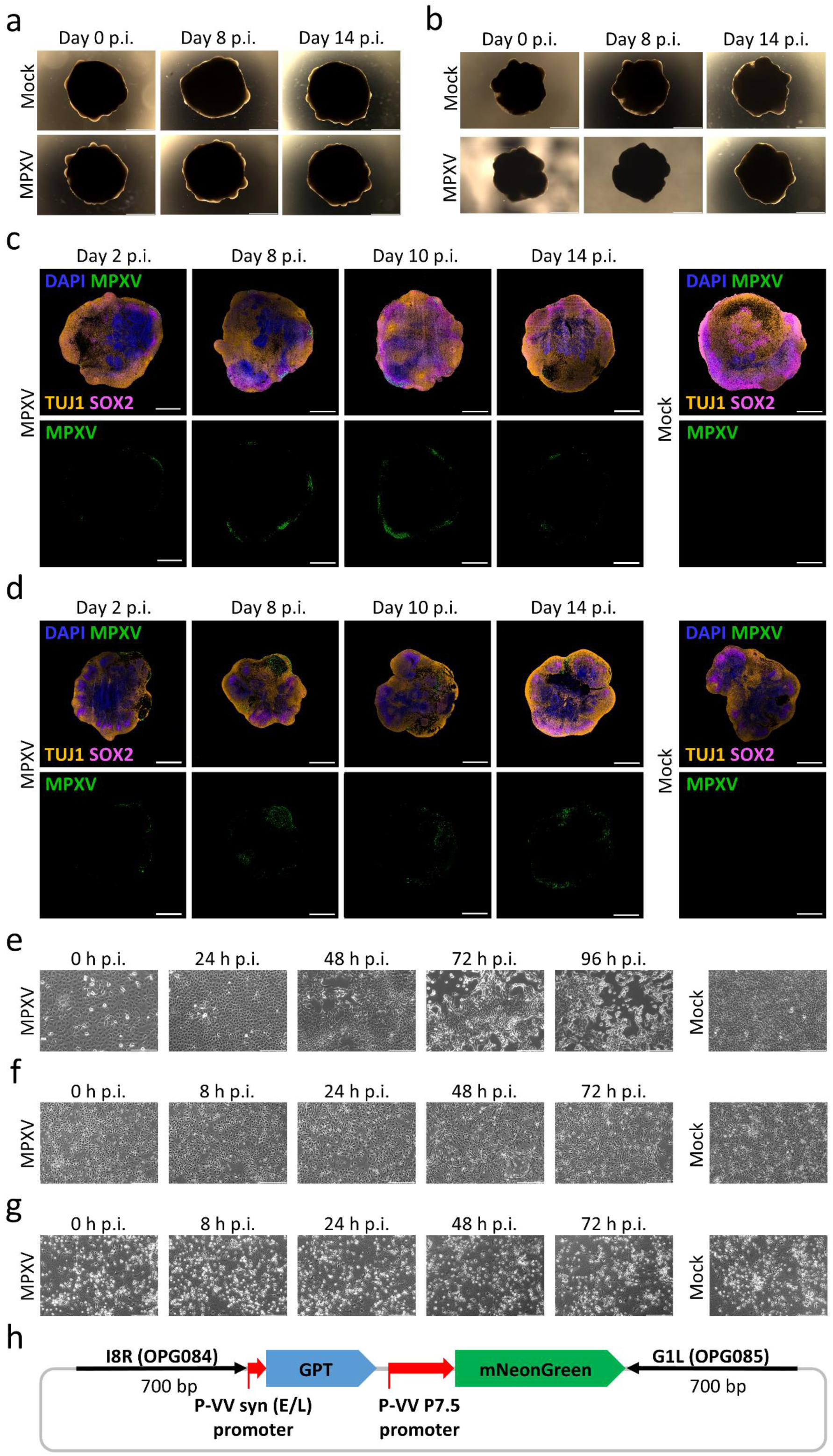
MPXV infects H1- and IMR90-derived hNOs and induces limited cytopathic effect in 2D neural cultures. (a-b) Representative light microscopy images of mock-treated and MPXV-infected H1- (a) and IMR90- (b) derived organoids at defined time points p.i. Organoids were infected with a clade IIb lineage MPXV isolate at a rate of 0.1 TCID_50_/cell at 84 (a) and 75 (b) days of development. Scale bar, 2000 μm. (c-d) Representative micrographs of MPXV-challenged and mock H1- (c) and IMR90- (d) derived hNOs at selected time points p.i. Organoids were exposed to a clade IIb lineage MPXV isolate at a rate of 0.1 TCID_50_/cell at 84 (c) and 75 (d) days of development. Scale bar, 1000 μm. DAPI, blue; MPXV, green; TUJ1, orange; SOX2, pink. Scale bar, 1000 μm. (e-f) Representative light microscopy images of 2D mock-treated and MXPV-infected Vero E6 (e), NPC (f) and neuron cultures (g). 2D cultures were infected with a clade IIb lineage MPXV isolate at a rate of 0.01 TCID_50_/cell. Scale bar, 200 μm. (h) Map of the pUC57-MPXV-GPT-mNG transfer vector (see “Material and methods”).

**Supplementary Figure 4:**
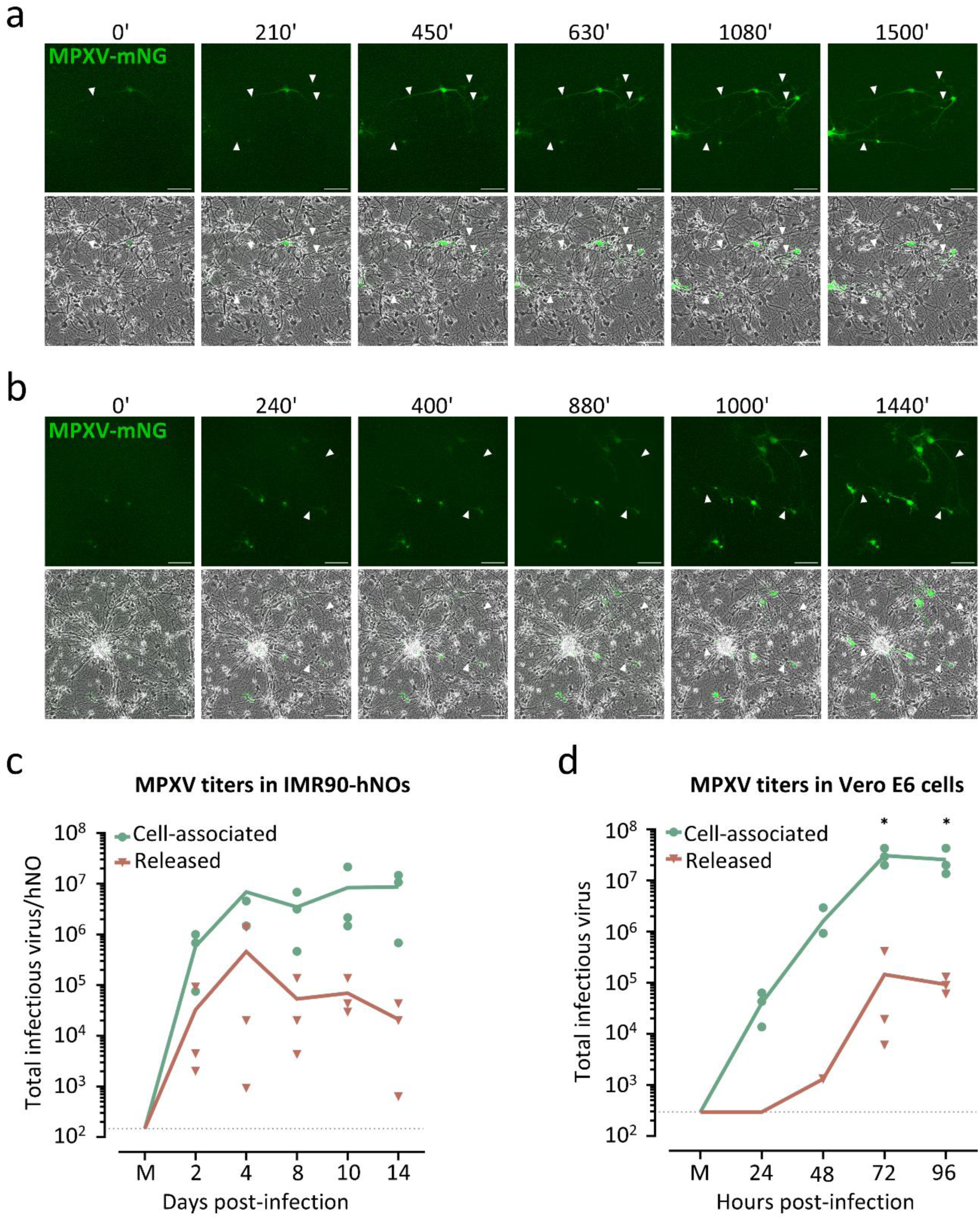
MPXV exploits neurite-mediated transport in 2D neurons and shows cell-associated spread in IMR90-derived hNOs and Vero E6 cells. (a-b) Representative images of 2D cultured neurons exposed to an mNG-expressing clade IIb lineage MPXV reporter strain at a rate of 0.01 TCID_50_/cell, showing viral spread between interconnected neighboring cells through neurite-mediated transport. White arrowheads indicate mNG-harboring neurites. Images were taken at 30 (a) and 40 (b)-minute intervals, time represented in minutes. Scale bar, 100 µm. (c) Cell-associated and released infectious MPXV 2 to 14 days p.i. of 75-86 days old IMR90-derived hNOs infected with a clade IIb lineage isolate at a rate of 0.1 TCID_50_/cell. Each symbol represents an individual organoid (n = 3 independent organoid batches). Unpaired student’s t-test was applied to compare groups. *p < 0.05, **p < 0.01. (d) Cell-associated and released MPXV titers 24 to 96 hours p.i. in Vero E6 cells challenged with a clade IIb lineage MPXV isolate at a rate of 0.01 TCID_50_/cell. Each symbol represents an individual Vero E6 batch (n = 3 independent Vero E6 batches). Unpaired student’s t-test was applied to compare groups. *p < 0.05, **p < 0.01.

**Supplementary Figure 5:**
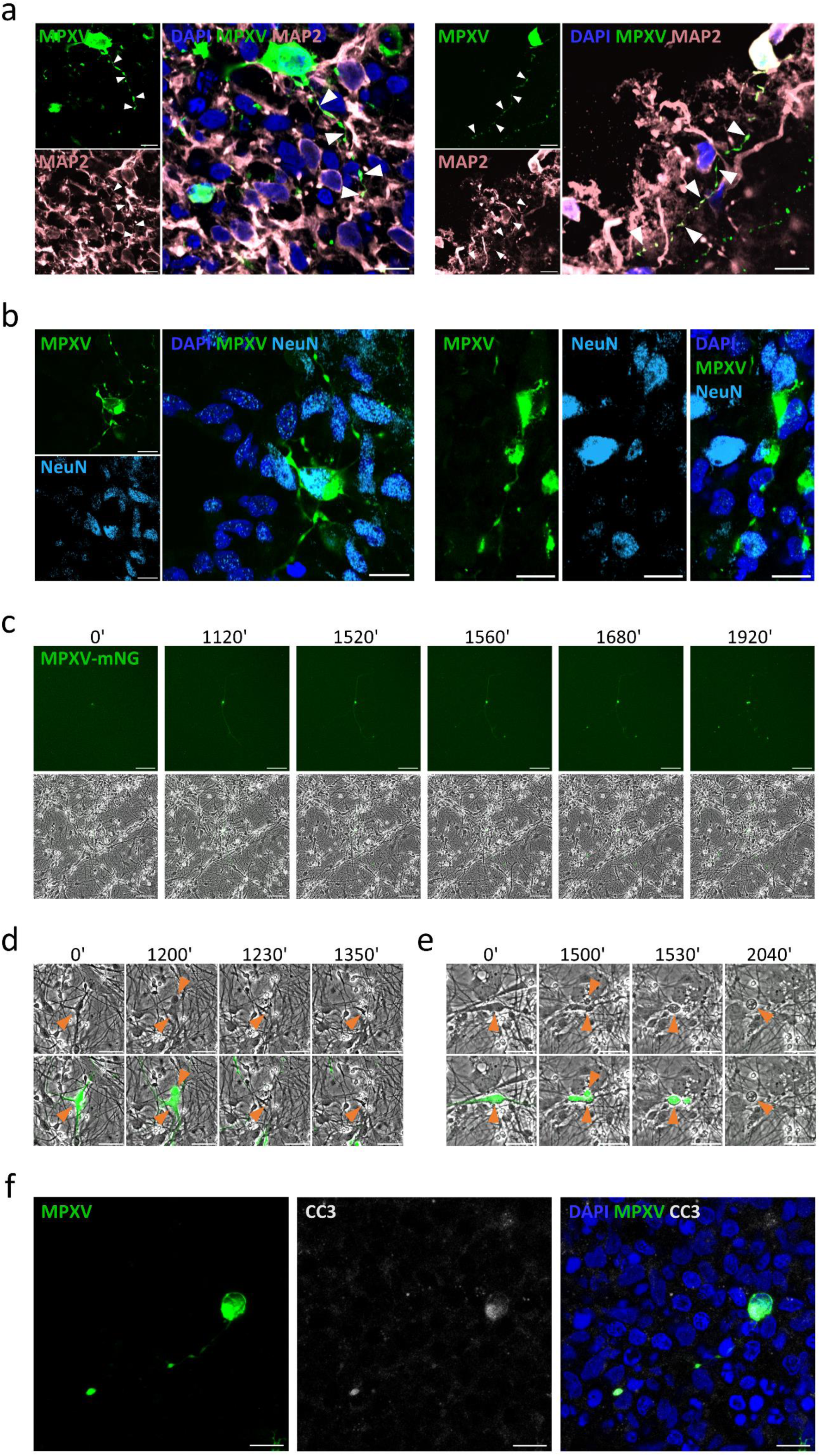
Neuritic beading preludes cell death in 2D cultured neurons. (a-b) Representative images of beaded MPXV-antigen harboring filaments co-localizing with alternative neuronal markers MAP2 and NeuN in both filaments and beads. White arrowheads indicate maker co-localization within beaded filament regions. DAPI, dark blue; MPXV, green; MAP2, mauve pink; NeuN, light blue. Scale bar, 10 µm. (c) Representative micrographs of 2D forebrain neurons infected with an mNG-expressing clade IIb lineage MPXV reporter strain displaying neuritic beading on filaments of a virus-harboring cell preceding mNG signal loss in neurites and cell rounding. Images were taken at 40-minute intervals, time represented in minutes. Scale bar, 100 µm. (d-e) Exemplary micrographs of documented changes in cell morphology in 2D neurons challenged with a MPXV clade IIb lineage mNG reporter strain. Morphological alterations were observed preceding and following neuritic beading, and included membrane blebbing, cell shrinkage (d), cell rounding and detachment (e). Orange arrowheads indicate cell bodies and membrane blebs. Images were taken at 30-minute intervals, time represented in minutes. Scale bar, 50 µm. (f) Representative micrographs of neuritic beads and soma of an infected cell accumulating CC3. DAPI, blue; MPXV, green; CC3, white. Scale bar, 10 μm.

**Supplementary Figure 6:**
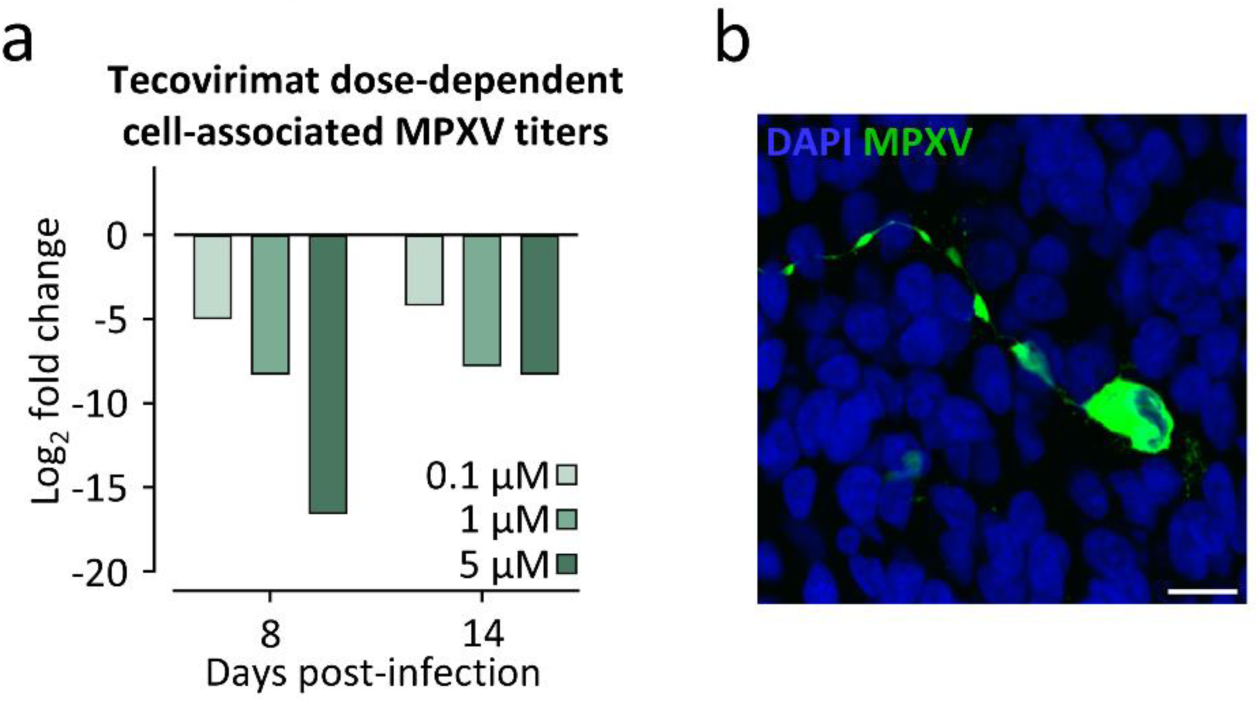
Tecovirimat treatment leads to dose-dependent reduction of cell-associated infectious MPXV but does not prevent neuritic beading. (a) Graph representing Log_2_ fold change of cell-associated live virus titers in H1-derived hNOs following MPXV challenge and tecovirimat treatment. H1-derived organoids of 110 days of development were infected with a clade IIb lineage MPXV isolate at a rate of 0.1 TCID_50_/cell. Medium supplemented with defined concentrations of tecovirimat was added 2 hours after initial MPXV exposure. n = 1 individual organoid batch. (b) Representative micrograph of cells displaying neuritic beading in tecovirimat-treated hNOs at 84 days of culture. DAPI, blue; MPXV, green. Scale bar, 10 μm.

**Supplementary Table 1:** DEGs in MPXV-infected versus mock-treated hNOs at day 4 p.i.

**Supplementary Table 2:** Overview of hNO batches and numbers per experiment.

**Supplementary Movie 1: MPXV-mNG infected NPCs show rosette-like virus spread**

**Supplementary Movie 2: MPXV-mNG infected neurons show somata and neurite infection**

**Supplementary Movie 3: Neurite-mediated MPXV-mNG spread – example 1**

**Supplementary Movie 4: Neurite-mediated MPXV-mNG spread – example 2**

**Supplementary Movie 5: Neuritic beading precedes neuron death in MPXV-mNG infected neurons – example 1**

**Supplementary Movie 6: Neuritic beading precedes neuron death in MPXV-mNG infected neurons – example 2**

